# Rfx6 promotes the differentiation of peptide-secreting enteroendocrine cells while repressing genetic programs controlling serotonin production

**DOI:** 10.1101/704924

**Authors:** Julie Piccand, Constance Vagne, Florence Blot, Aline Meunier, Anthony Beucher, Perrine Strasser, Mari L. Lund, Sabitri Ghimire, Laure Nivlet, Céline Lapp, Natalia Petersen, Maja S. Engelstoft, Christelle Thibault-Carpentier, Céline Keime, Sara Jimenez Correa, Valérie Schreiber, Nacho Molina, Thue W. Schwartz, Adèle De Arcangelis, Gérard Gradwohl

**Affiliations:** Institut de Génétique et de Biologie Moléculaire et Cellulaire (IGBMC), 1 rue Laurent Fries, 67404 Illkirch, France; Institut National de la Santé et de la Recherche Médicale (INSERM) U1258, 1 rue Laurent Fries, 67404 Illkirch, France; Centre National de Recherche Scientifique (CNRS) UMR7104, 1 rue Laurent Fries, 67404 Illkirch, France; Université de Strasbourg, 1 rue Laurent Fries, 67404 Illkirch, France; Centre for Metabolic Receptology, Novo Nordisk Foundation Center for Basic Metabolic Research, Faculty of Health Science, University of Copenhagen, Denmark

**Keywords:** Rfx6, Neurog3, Lmx1a, intestine, cell fate, serotonin, enteroendocrine cells, enterochromaffin cells, malabsorption, Mitchell Riley Syndrome

## Abstract

**Objective:** Enteroendocrine cells (EECs) of the gastro-intestinal tract sense gut luminal factors and release peptide hormones or serotonin (5-HT) to coordinate energy uptake and storage. Our goal is to decipher the gene regulatory networks controlling EECs specification from enteroendocrine progenitors. In this context, we studied the role of the transcription factor Rfx6 which had been identified as the cause of Mitchell-Riley syndrome characterized by neonatal diabetes and congenital malabsorptive diarrhea. We previously reported that Rfx6 was essential for pancreatic beta cell development and function, however, the role of Rfx6 in EECs differentiation remained to be elucidated.

**Methods:** We examined the molecular, cellular and metabolic consequences of constitutive and conditional deletion of Rfx6 in the embryonic and adult mouse intestine. We performed single cell and bulk RNA-Seq to characterize EECs diversity and identify Rfx6-regulated genes.

**Results:** Rfx6 is expressed in the gut endoderm; later it is turned on in, and restricted to, enteroendocrine progenitors and persists in hormone-positive EECs. In the embryonic intestine, the constitutive lack of Rfx6 leads to gastric heterotopia, suggesting a role in the maintenance of intestinal identity. In the absence of intestinal Rfx6, EECs differentiation is severely impaired both in the embryo and adult. However, the number of serotonin-producing enterochromaffin cells and mucosal 5-HT content are increased. Concomitantly, Neurog3-positive enteroendocrine progenitors accumulate. Combined analysis of single-cell and bulk RNA-Seq data revealed that enteroendocrine progenitors differentiate in two main cell trajectories, the enterochromaffin (EC) cells and the Peptidergic Enteroendocrine (PE) cells, whose differentiation programs are differentially regulated by Rfx6. Rfx6 operates upstream of *Arx*, *Pax6* and *Isl1* to trigger the differentiation of peptidergic EECs such as GIP-, GLP-1- or CCK-secreting cells. On the contrary, Rfx6 represses *Lmx1a* and *Tph1*, two genes essential for serotonin biosynthesis. Finally, we identified transcriptional changes uncovering adaptive responses to the prolonged lack of enteroendocrine hormones and leading to malabsorption and lower food efficiency ratio in Rfx6-deficient mouse intestine.

**Conclusion:** These studies identify Rfx6 as an essential transcriptional regulator of EECs specification and shed light on the molecular mechanisms of intestinal failures in human RFX6-deficiencies such as Mitchell-Riley syndrome.

## 1. INTRODUCTION

Enteroendocrine cells (EECs) are one of the five main intestinal cell subtypes, which include enterocytes, goblet-, Paneth- and tuft-cells. EECs are scattered as individual cells in the lining of the gut epithelium from the duodenum to the colon. Although they represent 1% of all intestinal epithelial cells they constitute the largest endocrine system of the body. EECs sense gut luminal factors (nutrients, microbial components) that trigger the secretion of peptide hormones or amines regulating food intake, digestion or glucose metabolism. Peptide hormones include Glucagon-Like Peptide 1 and 2 (GLP-1, GLP-2), Peptide YY (PYY), Gastric Inhibitory Peptide (GIP), Somatostatin (SST), Cholecystokinin (CCK), Secretin (SCT), Gastrin (GAST), Neurotensin (NTS) and Ghrelin (GHRL). The monoamine serotonin (5-hydroxytryptamine; 5-HT) is produced by enterochromaffin cells, the most abundant enteroendocrine cell type. Historically, EECs were classified using a one letter code based on their location and main secretory product. However, co-expression [1, 2] and single-cell transcriptomic studies [3, 4] tend to support the obsolescence of this classification [5] given that virtually all EECs express several hormones albeit not at the same level. Recent findings suggest that the hormonal repertoire of EECs might be modulated, along the crypt-villi axis, by local signals [6, 7]. Despite these major breakthroughs, our understanding of the molecular mechanisms controlling the differentiation of EECs is still limited, both during development and renewal in the adult intestine.

Several transcription factors have been demonstrated to regulate EECs differentiation. A master regulator is the basic helix-loop-helix (bHLH) transcription factor Neurogenin3 (Ngn3 or Neurog3) which is expressed in endocrine progenitors and determines endocrine commitment in the embryonic [8–10] and adult intestine [11]. Neurog3-expressing cells are rare hormone–negative endocrine progenitors confined to the crypts and Neurog3 expression is turned off when cells migrate towards the villi and differentiate into hormone-expressing cells. Indeed, *Neurog3*-deficient mice (constitutive and intestinal deletion) lack EECs. The loss of all enteroendocrine cells in mice leads to growth retardation, impaired lipid absorption and increased lethality, underlying the importance of enteroendocrine function [11]. Neurog3-positive endocrine progenitors arise from intermediate crypt progenitors which express the bHLH transcription factor Atoh1 [12]. Downstream of Neurog3, a complex network of transcription factors control the differentiation of subpopulations of EECs such as NeuroD1 [13], Arx [14, 15], Pax4 [14, 16], Isl1 [17], Pax6 [16], Foxa1/2 [18] or Insm1 [19].

Transcription factors controlling EECs differentiation, in almost all cases, also regulate pancreatic islet cell differentiation, suggesting shared genetic programs. We and others have reported the critical role of the transcription factor Regulatory Factor X 6 (Rfx6) in beta-cell development [20, 21] and function [22]. A previous study reported the regulation of *Gip* by Rfx6 in STC1 cell line [23]. Still, Rfx6 expression and role in mouse intestine and EECs differentiation is unknown. Rfx6, initially expressed broadly in the gut endoderm, becomes restricted to the endocrine lineage in the pancreas during embryogenesis and persists in adult islet cells. Rfx6 is necessary for proper islet cell development in zebrafish, mouse and Xenopus [20, 21, 24]. Rfx6-deficient mice lack alpha-, beta- and delta-cells, they are diabetic and die shortly after birth. Rfx6 loss in adult beta cells leads to glucose intolerance, impaired beta-cell glucose sensing, and defective insulin secretion [22]. In humans, several mutations in *RFX6* were identified as the cause of an autosomal recessive syndrome, named Mitchell-Riley syndrome, characterized by neonatal or childhood diabetes comprising hepatobiliary abnormalities and intestinal atresia [21, 25–32]. Most patients are presented with severe congenital malabsorptive diarrhea suggesting impaired EECs differentiation which however was not studied extensively.

In this study wa have investigated the function of Rfx6 in EECs differentiation in the embryonic and adult mouse. We have shown that EECs differentiation is severely impaired in *Rfx6*-deficient embryos, and only enterochromaffin cells are spared. Furthermore, we have detected patches of gastric cells in the small intestine of mutant newborns indicating gastric heterotopia. Constitutive intestinal deletion of *Rfx6* is found to be lethal at early post-natal stages. Deletion of *Rfx6* in the adult intestine is found to induces diarrhea, impaired lipid absorption and impaired food efficiency. Like in the embryo, EECs expressing peptide hormones were either happened to be lost or decreased in representation, while serotonin-positive enterochromaffin cells still developed with even slight increase in their number. Concomitantly, an increased number of Neurog3-positive enteroendocrine progenitors was also observed. Contrary to *in vivo* data, the removal of *Rfx6* in small intestinal organoids was found to result in impaired differentiation of all EECs, including enterochromaffin cells. By comparative transcriptomic studies, we have determined early Rfx6-dependent targets in the EEC lineage, and have identified secondary enhanced expression of neoglucogenic and nutrient absorption machinery genes reflecting adaptive response to the absence of enteroendocrine hormones. In parallel single-cell transcriptomic studies of EECs, we have described the dynamics of *Rfx6* expression and expression of other known and novel intestinal transcription factors. Overall, our results have shown that enteroendocrine progenitors differentiate in two main cell trajectories, the enterochromaffin (EC) cells and the Peptidergic Enteroendocrine (PE) cells, whose differentiation programs are differentially regulated by Rfx6.

## 2. MATERIAL AND METHODS

### 2.1 Animals and animal handling

All mice were housed in an animal facility licensed by the French Ministry of Agriculture (Agreement no. B67-218-5) and all animal experiments were supervised by GG (agreement no. C67-59) and approved by the Direction des Services Vétérinaires in compliance with the European legislation on care and use of laboratory animals. Rfx6^fl/+^ mice have been described previously [33] and were maintained on a heterozygous animals with C57BL/6N (Taconic) background. Rfx6^+/-^ mice have been generated by crossing Rfx6^f/+^ females with CMV-Cre males. Neurog3-Cre mice are a gift from Dr. Shosei Yoshida [34], Villin-Cre and Villin-CreER^T2^ were generously given by Dr. Sylvie Robine [35]. Neurog3^eYFP/+^ mice have been described previously [36]. Genomic tail DNA was analyzed by PCR using the primers detailed below. To achieve recombination in inducible mutants, the mice were treated with tamoxifen (10mg) (Sigma) by gavage twice per day, every second day during 5 days. Primers for genotypic were as follows:

*Neurog3* Forward: ctgcagtttagcagaacttcagaggga

*Cre* Reverse: atcaacgttttgttttcgga

*Villin* foward: ataggaagccagtttcccttc

*ERT* forward: gcattaccggtcgatgcaacgagtgatgag

*ERT* reverse: aggatctctagccaggcaca

*Rfx6* forward: gaaggtgcacccataaaagc

*Rfx6* reverse: tataagccacccagggtcag

*Neurog3* forward: cggcagatttgaatgagggc

*Neurog3* reverse: tctcgcctcttctggctttc

*GFP* forward: cctgaagttcatctgcaccac

*GFP* reverse: ttgtagttgtactccagcttgtgc

### 2.2 Histopathology and immunohistochemistry

Mouse tissues were fixed in 4% paraformaldehyde at 4°C overnight and embedded in paraffin or Sandon Cryomatrix (Thermo Scientific). Standard histology techniques were used: - for conventional histology, 7 µm paraffin sections were stained with Harris haematoxylin and eosin (H&E); - for goblet cells analysis, 7 µm paraffin sections were stained with Periodic Acid-Schiff (PAS) and haematoxylin or Alcian blue (AB) (pH 2.5); - for lipid and fat deposits detection, 10 µm cryo-sections were stained with Oil red O. Immunostaining on 7 (paraffin) or 10 (cryo) μm sections was performed using standard protocols. For BrdU detection assays, BrdU was injected intraperitoneally at 100mg/kg body weight, 24h prior sacrifice. If required, antigen retrieval was performed in 10mM Sodium Citrate pH6 in a pressure cooker. Primary antibodies used: guinea pig anti-Neurog3 1:1000 (M. Sander, UCSD, CA, USA); goat anti-Neurog3 1:2000 (G. Gu, Vanderbilt University, TN, USA); rabbit anti-Neurog3 1:500 (2763; IGBMC); rat anti-somatostatin 1:500 (Chemicon); mouse anti-ghrelin (C. Tomasetto, IGBMC, Strasbourg, France); goat anti-chromograninA 1:200 (Santa Cruz); goat anti-Cck 1:50 (Santa Cruz); guinea pig anti-glucagon 1:2000 (Linco); goat anti-glp1 1:100 (Santa Cruz); goat anti-Gip 1:100 (Santa Cruz); goat anti-gastrin 1:50 (Santa Cruz); goat anti-secretin 1:50 (Santa Cruz); goat anti-serotonin 1:2000 (Abcam); rabbit anti-serotonin 1:2000 (Diasorin Incstar); rat anti-BrdU 1:10 (AbD Serotec), rabbit anti-Rfx6 1:500 (2766, IGBMC); chicken anti-GFP 1:2000 (Abcam). Secondary antibodies conjugated to Alexa Fluor® 594, DyLight® 488, DyLight® 549 and DyLight® 649 (Jackson ImmunoResearch) were used at 1:500. For BrdU, Neurog3 and Rfx6, signal amplification was performed using biotin anti-rabbit (JacksonImmunoResearch) coupled antibody at 1:500 and streptavidin-Cy3 (or-488) conjugate at 1:500 (Molecular Probes). Immunofluorescent stainings on mouse intestinal organoids were realized as previously described (O’Rourke K., P. et al., 2016).

### 2.3 Generation and maintenance and 4-hydroxytamoxifen treatment of mouse intestinal organoids

Mouse intestinal organoids were generated as previously described (Sato et al., 2009) from small intestine of 2-6 month-old CD1 Neurog3^eYFP/+^ mice, Rfx6^fl/fl^ and Rfx6^fl/fl^; Villin CreER^T2^ mice. Briefly, crypts from duodenum or jejunum were isolated and cultured into Matrigel drops (Fisher Scientific^TM^) to develop organoids in presence of the following medium: Advanced DMEM/F-12, HEPES 10 mM, GlutaMax 2 mM, penicillin/streptomycin 100 U/mL, supplement B27 1X (all from Gibco^TM^), N-acetyl-L-cysteine 1 µM (Merck), human recombinant R-spondin-1 500 ng/mL (PeproTech^®^), murine recombinant EGF 50 ng/mL (PeproTech^®^) and human recombinant Noggin 100 ng/mL (R&D Systems^®^). Organoids were maintained at 37°C, 5% CO2 and were split into 24-well plates every 5 to 7 days. To induce *in vitro* Cre recombinase activation and specific deletion of *Rfx6*, organoids derived from Rfx6^fl/fl^ (controls) and Rfx6^fl/fl^; Villin-CreER^T2^ duodenum (n=4 per genotype) were treated two days after seeding with 4-hydroxytamoxifen (4-OHT; Sigma-Aldrich) 1.25 µM for 15h, after which the medium was replaced with fresh medium without 4-OHT. Organoids were harvested 8 days after 4-OHT treatment for RNA extraction.

### 2.4 Measurement of intestinal serotonin content

After 11 days of tamoxifen injections, four adult male Rfx6^fl/fl^ and three Rfx6^fl/fl^ Villin-CreER^T2^ were euthanized by cervical dislocation and the entire gastrointestinal tract were excised. Using dermal biopsy punches (1.5 mm) intestinal samples were obtained from relevant sites throughout the GI tract (Duodenum, Jejunum, Ileum, Proximal Colon and Distal Colon). All samples were quickly homogenized and purified in ice-cold 1M perchloric acid to precipitate proteins. Following centrifugation at 14.000G at 4°C, 15 min, where after the supernatant were collected and taken through an additional purifying step adding similar volume 1M perchloric acid followed by mixing and centrifugation. The protein-free samples were diluted 2 or 10 times in MilliQ water prior HPLC injection (10µl). 5-HT concentration was measured by HPLC-ECD (HTEC-500, Eicom) with a PP-ODS2 column (Amuza) following manufacturer’s instructions regarding flow rate, mobile phase and applied potential. Retention time (3.8 seconds) for 5-HT were determined by a control 5-HT dilution of 1pM (Sigma-Aldrich). Results are stated as area under the curve (AUC) and presented as mean ± SD. Unpaired T-test was applied to calculate significance (*p≤0.05; **p≤0.01).

### 2.5 Morphometric analysis

Neurog3-positive cells or Neurog3/BrdU double positive cells were counted in adult mice after immunostaining on approximately 50 sections of duodenum, jejunum, ileum and colon three months after tamoxifen treatment (n=3 per genotype). The number of Neurog3-positive cells was normalized according to the area of the sections estimated by the surface of DAPI staining. The number of Neurog3/BrdU double positive cells was normalized to the total number of Neurog3 cells, and expressed as a percentage. 5-HT positive cells were counted after immunostaining on sections of duodenum and ileum from adult animals (n=3 per genotype), one week after tamoxifen treatment. At least 50 well-oriented crypt-villus sections per animal were analyzed and as indicated above the number of 5-HT-positive cells was normalized according to the DAPI-positive area.

### 2.6 Metabolic studies

Mice were fed with normal chow diet (DO3, SAFE) from weaning. Glucose tolerance tests were performed as described previously [22]. Food intake and feces production were measured over a 48h period. Feces energy content, energy excretion, energy ingested and food efficiency were measured by the Mouse Clinical Institute (MCI) metabolic platform (http://www.ics-mci.fr). Blood analysis were performed on plasma from adult males by the MCI. The energy content of the stools was evaluated using a bomb calorimeter (C503 control, IKA) The energy excreted is calculated as Feces Energy Content (Cal/g) X Feces Weight (g) / 2. The energy ingested is calculated as food intake per day (g) X diet calorific value (3.5 Kcal/g). The energy digested is calculated by the difference between total calories ingested and excreted in feces. It is then expressed as “food efficiency”.

### 2.7 Statistics

Comparisons of means and medians were performed with Graphpad Prism 7.0 or R softwares. Values are presented as mean ± SD or SEM. p-values were determined using the 2-tailed Student t-test with unequal variance or the 2-tailed nonparametric test of Mann-Whitney offered in Prism. p≤ 0.05 was accepted as statistically significant.

### 2.8 Dissociation of mouse intestinal organoids into single cells

Organoids were dissociated to single cells as previously described, with some modifications (Grün et al., Nature, 2015). Briefly, after medium removal and Matrigel disruption, organoids were dissociated in 500 µL of TrypLE^TM^ Express Enzyme (1X) (Gibco^TM^) at 37°C for 20 min, assisted by a mechanical dissociation (up and down with a p1000 and a final passage through a syringe fitted with a 18G needle). The dissociation was stopped in a medium containing Advanced DMEM/F-12, fetal calf serum 10%, N-acetylcysteine 0.5 mM, and supplement B27 1X. The cells were pelleted, washed 2 times and resuspended in the same medium. Finally, they were strained through a 50 µm cell strainer, and stained with DAPI (to exclude dead cells) before to be sorted by flow cytometry (FACS Aria II, BD).

### 2.9 RNA isolation and qPCR

Total RNA from whole small intestine (pups), duodenum (adults), jejunum (adults), ileum (adults) or colon (pups and adults) was extracted using TRI Reagent (Invitrogen) or RNeasy® Midi Kit (Qiagen), according to the manufacturer’s instructions. Total RNA from organoids was extracted using RNeasy® Mini Kit (Qiagen) according to the manufacturer’s instructions. Reverse transcription was performed using Transcriptor Reverse Transcriptase (Roche) and Random primers (Roche). Quantitative PCRs were performed using mouse-specific TaqMan primers and probes (Applied Biosystems) recognizing Neurog3 (Mm00437606_s1), ChgA (Mm00514341_m1), Arx (Mm00545903_m1), Pax4 (Mm01159036_m1), Pax6 (Mm00443081_m1), Isl1 (Mm00517585_m1), Lmx1a (Mm00473947_m1), Tac1 (Mm01166996_m1), Ucn3 (Mm00453206_s1), NeuroD1 (Mm00520715_m1), Pyy (Mm00520715_m1), Nts (Mm00481140_m1), Cck (Mm00446170_m1), Sct (Mm00441235_g1), Gip (Mm00433601_m1), Gcg/Glp1 (Mm00801712_m1), Tph1 (Mm00493794_m1), Gast (Mm00772211_g1), Sst (Mm00436671_m1), Ghrl (Mm00445450_m1), Nkx2.2 (as described in Gross S. et al., 2016; Forward primer: cctccccgagtggcagat; Reverse primer: gagttctatcctctccaaaagttcaaa; FAM-MGB Probe: ccattgactctgccccatcgctct) or UPL probes #83 for Rfx6 (Forward primer: tgcaggaaagagaaactggag; Reverse primer: ggaaatttttggcgaattgtc), with Light Cycler 480 Probes Master Mix (Roche) on Light Cycler 480 (Roche). Gene expression levels were normalized to Rplp0 (Mm01974474_gH).

### 2.10 Microarrays

Samples processing: Microarray analysis was performed according to Agilent protocol “One-Color Microarray-Based Gene Expression Analysis – Low Input Quick Amp Labeling” version 6.5, May 2010 (Cat # G4140-90040). Complementary RNA (cRNA) samples were linearly amplified and labeled with cyanine 3 starting from 100 ng of total RNA. Following fragmentation, labeled cRNA were hybridized on Agilent “SurePrint G3 Mouse Gene Expression 8×60K Microarray” (Design ID: 028005), for 17 hrs, at 65°C under 10 rpm. Following washing, the slides were scanned using an Agilent G2565CA microarray Scanner System, at a 3 µm resolution in a 20-bit scan mode, according to the “AgilentG3_GX_1Color” protocol. *Data analysis:* Raw .tif images were then extracted using Agilent “Feature Extraction, version 10.10.1.1” following “GE1_1010_Sep10” protocol. Median raw expression values were further normalized by quantile method [37]. Differential expression analysis between Rfx6*^-/-^* vs controls and *Rfx6^ΔEndo^* vs controls) were performed using the R package limma version 3.36.5 [38]. Data have been deposited on GEO repository (GSE133038).

### 2.11 RNA Sequencing

Ileon from adult Rfx6^ΔAdInt^ mice was harvested at 8 days and 3 months after tamoxifen gavage. Total RNA was extracted using TRI-reagent or RNeasy® Midi Kit (Qiagen). Libraries were prepared using the total RNA-seq Ribo-zero protocol and sequenced on an Illumina HiSeq 2500 (single-end 50bp reads). Reads were identified and mapped onto the mm9 assembly of mouse genome using Tophat version 2.0.10 and the Bowtie2 version 2.1.0 aligner. Quantification of gene expression was performed using HTSeq version 0.6.1 and gene annotations from Ensembl release 67. Normalization of read counts and differential expression analysis between controls and Rfx6^ΔAdInt^ samples were performed using the method implemented in the DESeq2 Bioconductor library version 1.0.19 [39]. Data have been deposited on GEO repository (GSE133038).

### 2.12 Single-cell RNA-Seq; sample processing

eYFP^+^ cells, from dissociated Neurog3^eYFP/+^ intestinal organoids, were sorted directly into four 96 well plates « Precise WTA Single Cell Encoding Plate » (BD^TM^ Precise WTA Single Cell Kit, BD Genomics) (1 cell/well) using a FACSAria Fusion (BD). DAPI was used to sort only living cells. The library was prepared according manufacturer’s recommendations (910000014 Rev. 02 11/2016, BD Genomics) and sequenced on an Illumina HiSeq 4000 using paired-end 2×100bp lanes sequencing. Data have been deposited on GEO repository (GSE133038).

### 2.13 Single-cell RNA-Seq analysis

#### 2.13.1 Demultiplexing, alignment and quantification

Fastq files were demultiplexed using the sample index encoded in reads 1. Reads 2 having a bad quality (--quality-cutoff 20,20) or corresponding to polyA sequences or to adapters were removed using Cutadapt version 1.10. Reads 2 mapping to rRNA were identified using Bowtie 2 version 2.2.8 and removed. Reads 2 were aligned to mm10 genome using STAR version 2.5.3a. After alignment, reads corresponding to technical duplicates were removed using UMI-Tools version 0.4.4 (method: “directional-adjacency”) [40]. Quantification of gene expression was performed using HTSeq version 0.6.1.post1 (parameter “--mode union”) and gene annotations from Ensembl release 90.

#### 2.13.2 Quality filtering of cells

Cells were filtered in function of several quality criteria: (i) Percentage of reads uniquely aligned to genome had to be superior or equal to 40%, (ii) percentage of UMIs assigned to genes had to be superior or equal to 50%, (iii) number of assigned UMIs had to be superior or equal to 200,000, (iv) proportion of UMIs assigned to mitochondrial genes had to be inferior to 20%. Finally, 290 cells were kept for further analyses. Among these filtered cells, the number of assigned UMIs varies from 200,000 to 2.3 million. Regarding the number of expressed genes per cell, it varies from 2,777 to 10,019 with a median equal to 4,606.

#### 2.13.3 Normalization, dimensionality reduction and clustering

Data were then analyzed using the Seurat package version 2.3.4 [41]. Before creating the Seurat object, a gene filtering was performed. On the 29,161 genes expressed in the data, only 9,057 were retained as they met two criteria: they were coding genes or lincRNA and had at least 5 UMIs assigned in 5 samples. The normalization and scaling of the data were performed using the functions NormalizeData (method “LogNormalize”) and ScaleData implemented in Seurat.

Variable genes were identified using the FindVariableGenes function in the Seurat package: the 830 genes having a scaled dispersion superior or equal to 1 and a log-normalized average expression superior or equal to 0.2 were selected as variable genes. Then, a PCA (Principal Component Analysis) was done with the function RunPCA of Seurat, based on these 830 variable genes. The 13 first axes were used to create an UMAP, as they had a significant PCA score, according to the “Jack Straw” approach implemented in Seurat. The UMAP was performed with the function RunUMAP in Seurat (n_neighbors = 20 and min_dist =0.3). Finally, the clustering was performed based on the two first axis of the UMAP, using the SNN (shared nearest neighbor) modularity optimization algorithm implemented in the FindClusters function of Seurat. The following parameters were used: k.param = 30, resolution =0.9 and prune.SNN = 5/15. Eight groups of cells were found.

#### 2.13.4 Identification of markers of the groups

Before performing the differential expression analysis, genes were filtered. On the 9,057 already filtered genes, we kept only 8,644 genes as they had a normalized expression equal or superior at 1 in at least 1% of the cells. Differential expression analysis was done using the QLF (quasi-likelihood) approach implemented in the R package edgeR version 3.20.6 [42]. The detection rate was added as a covariate to the model as it improves the performance of the analysis [43]. To compare one group of cells to the others, we used the following contrasts: 1 for said group and −1/7 for the seven other groups. One gene was considered as the marker of a group if it was significantly over-expressed in this group with a FDR inferior or equal to 10^-10^ and if it was expressed in at least 20% of the cells of the said group.

#### 2.13.5 Cell trajectory

The log-transformed gene expression matrix of the 830 variable genes was used to generate diffusion map, using the function DiffusionMap (k = 15) implemented in destiny R package, version 2.12.0 [44].

## 3. RESULTS

### 3.1 Rfx6 is expressed in endocrine progenitors in the crypts and in all nascent hormone-expressing cells at the base of villi in the adult mouse intestine

To determine Rfx6 intestinal expression, we performed a series of immunofluorescence experiments in the embryo, as well as in adult CD1 mice (Figure 1). It was previously shown that Rfx6 is expressed broadly in the gut endoderm at E9 [20, 21]. From E13.5, Rfx6 becomes restricted to a small number of scattered cells in the embryonic intestine in a pattern of developing enteroendocrine cells (Figure 1A and data not shown). Accordingly, we found that Rfx6 is expressed in Neurog3-positive enteroendocrine progenitors during embryogenesis and in the adult small intestine. Indeed, we observed that Rfx6 is expressed in a subset of crypt based Neurog3-positive endocrine progenitor cells (40%; n=300; Figure 1B-D). The majority of Rfx6-positive-nuclei are found in the crypts (Suppl. Figure 1). Nevertheless, Rfx6 is maintained in hormone-expressing enteroendocrine cell types including incretin (GLP-1 and GIP) producing cells, as well as 5-HT enterochromaffin cells (Figure 1G-N, Suppl. Figure 1). We observed that cells positive for both Rfx6 and hormones show different intensity of Rfx6 staining along the crypt-villus axis. Rfx6 staining appears to be strongest in the crypts and in the lower part of villi and becomes weaker while enteroendocrine cells differentiate and migrate up toward the villus top (Figure 1J). Indeed, Chromogranin A (ChgA)-positive or 5-HT-positive but Rfx6-negative cells can be found at the villus top indicating gradual loss of Rfx6 as enteroendocrine cells differentiate and migrate up (Figure 1F, Suppl. Figure 1). In contrast, all GIP-expressing cells expressed Rfx6 (Suppl. Figure 1).

**Figure 1:**
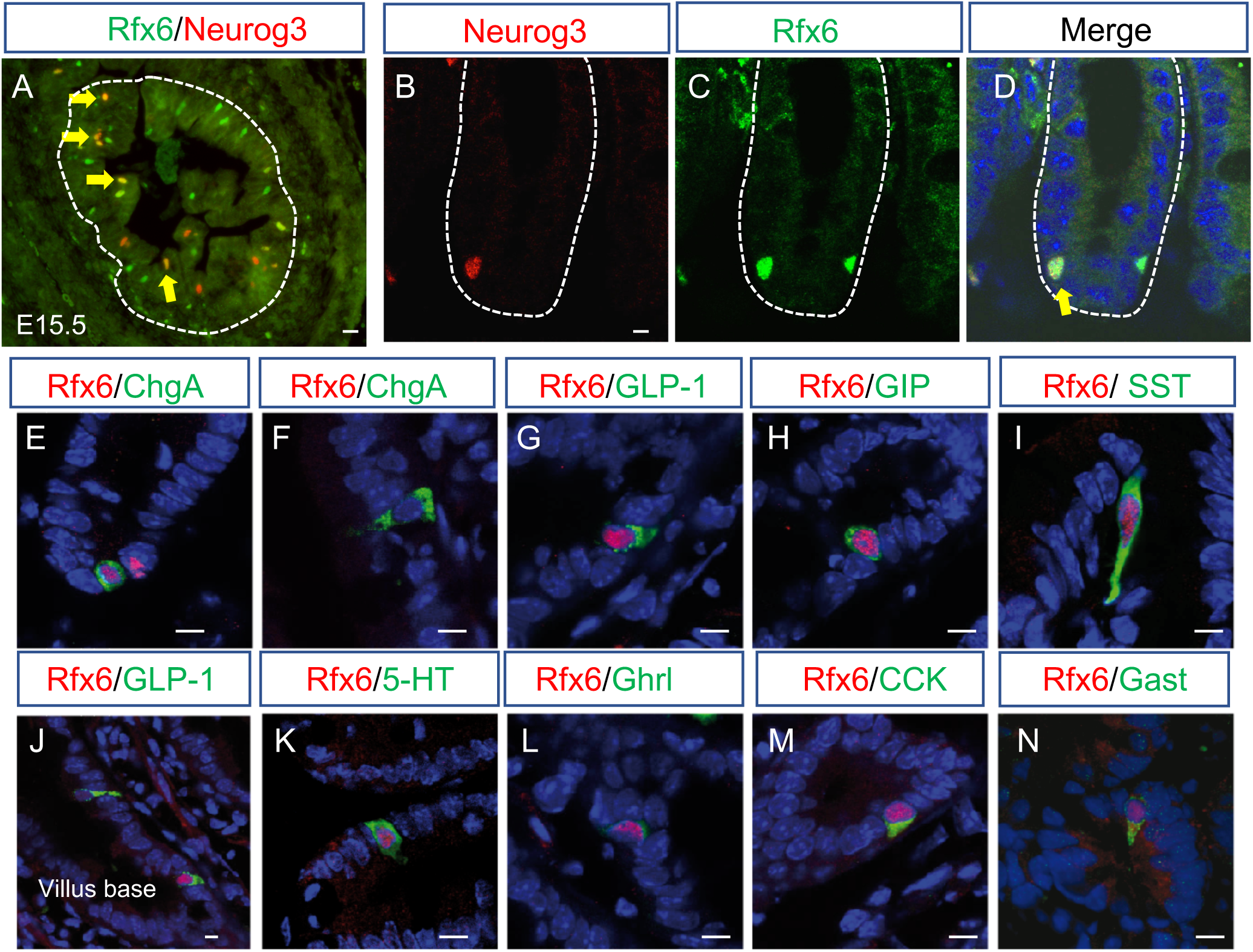
Expression of Rfx6 in the intestinal endocrine lineage of embryonic and adult mice. (A-D) Double immunofluorescence for Rfx6 (green) and Neurog3 (red) on cryosections of embryonic E15.5 (A) and adult small intestine (B-C). Yellow arrows point to Rfx6/Neurog3 double positive cells. (E-N) Double immunofluorescence for Rfx6 in red and ChromograninA (ChgA), Glucagon like peptide-1 (GLP-1), Gastric inhibitory Polypeptide (GIP), Somatostatin (Sst), Serotonin (5-HT), Ghrelin (Ghrl), Cholecystokinin (CCK) or Gastrin (Gast), in green. Nuclei are stained with DAPI (blue). Scale bar 10µM.

### 3.2 Ectopic expression of gastric genes and impaired enteroendocrine cell differentiation in the embryonic intestine lacking Rfx6

At first, Rfx6 is expressed in the embryonic gut endoderm and then becomes restricted to the enteroendocrine lineage at later stages of development suggesting different roles. To decipher Rfx6 intestinal functions, we compared the transcriptome of the embryonic small intestine of *Rfx6* null mice (*Rfx6^-/-^*), as well as of mice with an endocrine specific deletion of the gene *(Rfx6^f/f^; Neurog3*-Cre; called thereafter *Rfx6^ΔEndo^*) to wild-type controls. We observed a strong down-regulation of the vast majority of genes encoding enteroendocrine hormones, *Gcg (stands herein for preproglucagon gene)*, *Gip*, *Ghrl*, *Sst*, *Nts*, *Cck*, *Pyy*, *Sct*, both in *Rfx6^-/-^* as well as in *Rfx6^ΔEndo^* intestine (Figure 2 A, B, Table S1 and 2). Lmx1a, a transcription factor regulating *Tph1* expression and thus important for 5-HT production in EC cells, is up-regulated in *Rfx6-/-* intestine (Fold Change (FC) =1.7; adjusted p-value=0.01). Indeed, recent studies revealed the importance of Nkx2-2 transcription factor for enterochromaffin cell differentiation through control of *Lmx1a* expression [45]. Lmx1a in turn transactivates *Tph1* encoding Tryptophan hydroxylase 1, the rate-limiting enzyme for the production of peripheral serotonin. Accordingly, we observed a trend for *Tph1* to increase (FC= 1.3; p-value 0.016; adjusted p-value=0.13).

**Figure 2:**
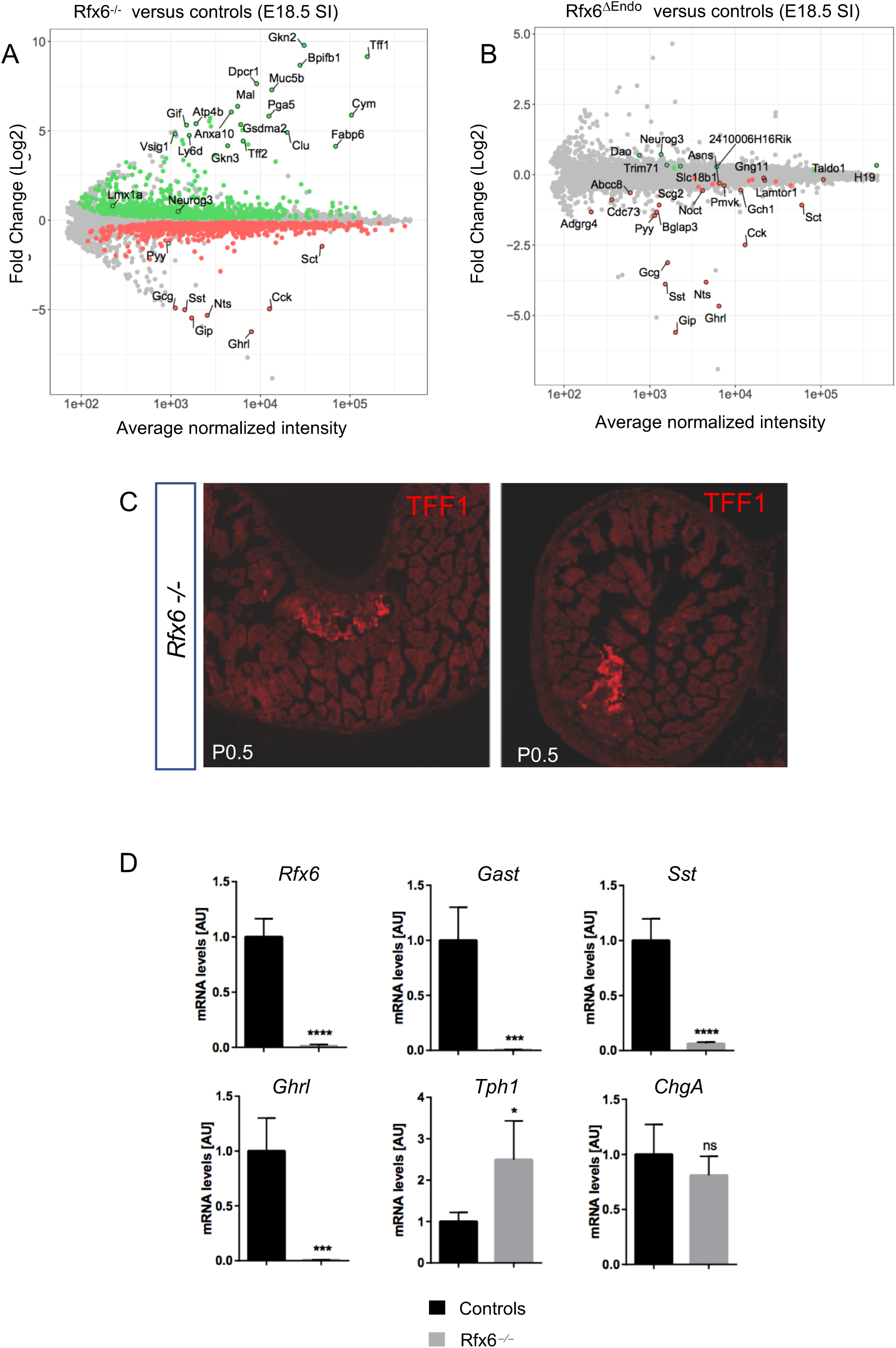
Severe impairment of enteroendocrine cell differentiation in the gastro-intestinal tract of *Rfx6*-deficient embryos and presence of gastric heterotopia in mutant neonates. (A, B) Gene expression profiling of the embryonic small intestine of mice with constitutive (*Rfx6^-/-^*) or endocrine-specific deletion (*Rfx6^ΔEndo^*) of Rfx6 at E18.5. (A) *Rfx6 ^-/-^* vs control comparison (micro-array data). MA-plot representing the log Fold-Change as a function of the mean of normalized intensity. Green and red points represent significantly (Adjusted p-value ≤ 0.1) up- and down-regulated genes, respectively. (B) *Rfx6^ΔEndo^* vs control comparison (micro-array data). MA-plot representing the log Fold-Change as a function of the mean of normalized intensity. Green and red points represent significantly (Adjusted p-value ≤ 0.1) up- and down-regulated genes, respectively. (C) Immunofluorescence for TFF1 on small intestine cryosections of two *Rfx6^-/-^* new born mice revealing gastric heterotopia. (D) RT-qPCR analysis of relevant genes in the stomach (P0.5) of *Rfx6^-/-^* and wild-type mice. Analysis was performed on n=4 animals per genotype. Data are represented as mean ± SD; unpaired t-test, **** p≤0.0001 ***, p≤0.001, ** p≤0.01, * p≤0.05, ns: not significant. AU stands for Arbitrary Unit. SI: Small Intestine. E18.5: Embryonic day 18.5. P0.5: postnatal day 0.5.

Unexpectedly, transcriptome comparison revealed that many gastric genes were strongly up-regulated in *Rfx6^-/-^* (Figure 2A), but not in *Rfx6^ΔEndo^* embryonic intestine, compared with wild-type. These genes include *Gkn2, Tff1* and *Muc5b,* which are normally expressed in pit cells of the gastric glands, as well as *Atp4b,* which encodes a subunit of the gastric proton pump expressed in parietal cells of the fundic gland. These data suggest that heterotopic gastric mucosa forms in the small intestine of *Rfx6*-deficient mice. We were able to confirm this hypothesis by RT-qPCR for *Gkn2* and *Tff1* (data not shown) and at the protein level for TFF1 (Figure 2C). In agreement with the transcriptomic data, we found many patches of TFF1-positive cells in the small intestine of *Rfx6*-deficient newborn mice but never in controls (not shown). Next, we investigated gastric enteroendocrine cell differentiation. *Tph1* expression was, similarly to the intestine, up-regulated, while the expression of *Gast*, *Sst* and *Ghrl* was severely decreased in the stomach of *Rfx6*-deficient newborn mice (Figure 2D). Thus, our study suggests that, in the mouse embryo, Rfx6 controls enteroendocrine cell differentiation in the gastro-intestinal tract and is required for the maintenance of intestinal cell identity.

### 3.3 Constitutive, but not adult-induced intestinal deletion of *Rfx6,* is lethal

To further study Rfx6 intestinal function and to bypass any effect related to its pancreatic function, we generated intestine-specific *Rfx6*-deficient mice (*Rfx6^fl/fl^*; *Villin-Cre*, referred as *Rfx6^ΔInt^*). *Rfx6^ΔInt^* mice died shortly after birth (between P2-P5, n=14 litters), except one animal who survived up to 13 weeks. The presence of milk in the stomach testified that mutants were fed properly, however, they sometimes failed to thrive. Similarly to *Rfx6^-/-^* embryonic intestine, RT-qPCRs revealed that *Gcg*, *Gip*, *Cck*, *Pyy*, *Nts*, *Sst* and *Ghrl* transcripts are mostly undetectable in newborns *Rfx6^ΔInt^* small intestine, *Sct* is decreased and *Tph1* increased (data not shown). Because of the early lethality of Rfx6^ΔInt^ pups, the role of Rfx6 in the adult intestine could not be studied in this model. We thus generated mice with an inducible deletion of *Rfx6* (*Rfx6^fl/fl^*; *Villin-CreER^T2^*, referred as *Rfx6^ΔAdInt^*). Tamoxifen gavage efficiently deleted *Rfx6,* as shown by the absence of wild-type *Rfx6* mRNA and protein in the small intestine and the colon of *Rfx6^ΔAdInt^* mice 8 days after treatment (Suppl. Figure 2A and data not shown). *Rfx6^ΔAdInt^* mice started to lose weight one week after tamoxifen treatment, however they caught up 7 days later (Figure 3A). Only one *Rfx6^ΔAdInt^* mice did not overcome the weight loss and died 11 days after the beginning of the treatment. Thus, except this single case, we did not observe any death upon Rfx6 intestinal deletion in the adult mouse (n≥100). In summary, the lack of functional intestinal Rfx6 is lethal at early postnatal stages but is not life threatening in the adult mice.

**Figure 3:**
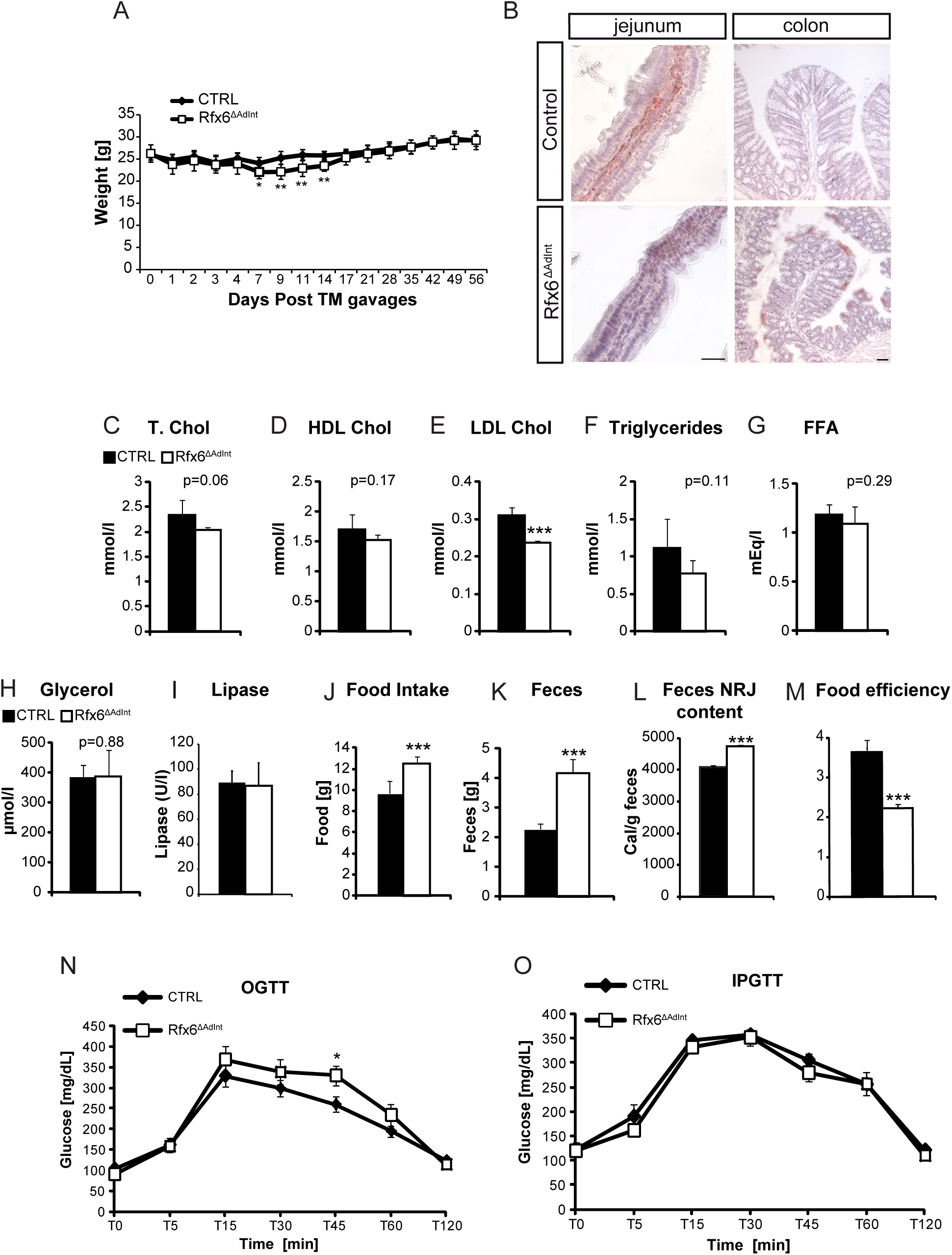
Energy homeostasis in *Rfx6^ΔAdInt^* mice. (A) Weight follow-up of controls and Rfx6^ΔAdInt^ mice after tamoxifen gavages. (B) Oil Red O staining on cryosections from jejunum and colon from Rfx6^ΔAdInt^ and control animals one month after tamoxifen gavage. (C-E) Blood concentration of total (T. Chol, C), high density lipoprotein (HDL Chol, D), and low-density lipoprotein cholesterol (LDL Chol, E) in Rfx6^ΔAdInt^ mice. (F-I) Blood concentration of triglycerides (F), free fatty acids (FFA) (G), glycerol (H) and lipase (I) in Rfx6^ΔAdInt^ mice. (J-M) Decreased food efficiency and increased food intake and feces production in Rfx6^ΔAdInt^ mice. Food intake (J) and feces production (K) of control and mutant mice during 48h. (L) Energy (NRJ) content of the stools was evaluated in a bomb calorimeter and Food Efficiency (M) calculated (calories ingested/excreted in feces). (C-F) analysis performed 3 months after tamoxifen treatment. (N, O) oral (N, OGTT) and intraperitoneal (O, IPGTT) glucose tolerance tests on adult control and Rfx6^ΔAdInt^ mice one month after tamoxifen-induced Rfx6 deletion. n=5-6 animals were analyzed per group. Data are represented as mean ± SEM; unpaired t-test, *** p≤0.001, ** p≤0.01*, p≤0.05.

### 3.4 Deletion of *Rfx6* in the adult intestine induces diarrhea and impairs lipid absorption and food efficiency

We next investigated the metabolic consequences of induced intestinal deletion of *Rfx6* in the adult mice one month after tamoxifen treatment (Figure 3). First, we noticed that mutant mice exhibited liquid and yellowish stools (data not shown) suggesting diarrhea and steatorrhea. In agreement with the later hypothesis, Oil red O staining of the mutant gut revealed a clear reduction in the presence of neutral lipids content in the enterocytes and the lamina propria in the small intestine as well as accumulation of lipid droplets in the colon (Figure 3B). In addition, although total cholesterol and HDL levels are not affected, levels of LDL cholesterol were reduced in the serum of *Rfx6^ΔAdInt^* mice (Figure 3 C-E). Triglycerides concentrations were also reduced, but not significantly (p=0.11), free fatty acids and glycerol levels were not affected (Figure 3F-H). Since serum pancreatic lipase levels are unchanged (Figure 3I), these results suggest that the reduced numbers of Oil red O-positive lipid droplets and the reduced serum levels of LDL cholesterol found in *Rfx6^ΔAdInt^* mice are due to impaired lipid absorption rather than impaired lipid processing.

Glucose clearance was explored by oral (OGTT) and intraperitoneal (IPGTT) glucose tolerance tests (Figure 3N, O). After glucose gavage, blood glucose concentration raised similarly in mutant and control mice suggesting normal intestinal glucose absorption. OGTT revealed that *Rfx6^ΔAdInt^* mice are slightly glucose intolerant. This effect could result from the lack of incretin hormones. Indeed, when the incretin effect is bypassed in IPGTT, glucose clearance is unchanged. To further characterize these mice, we performed analyses of food efficiency (Figure 3 J-M). Mice, in which *Rfx6* was deleted from the adult intestine, showed an increased production of feces due, at least in part, to an increased food intake (Figure 3J-K). Fecal energy measurement, using bomb calorimetry (Figure 3L), revealed fecal energy loss in *Rfx6^ΔAdInt^* mice compared to the controls, suggesting malabsorption. Despite increased food intake, *Rfx6^ΔAdInt^* mice have a decreased food efficiency ratio (Figure 3M). Together, these data suggest that decreased food efficiency of *Rfx6^ΔAdInt^* mice could result from diarrhea and lipid malabsorption. This is in agreement with previous work in mice lacking all enteroendocrine cells [11] as well as with symptoms of Mitchell Riley patients [21].

### 3.5 Impaired production of all enteroendocrine hormones except serotonin which is increased in *Rfx6*-deficient adult intestine

To decipher the mechanisms underlying Rfx6 function in the adult intestine we determined the genes differentially expressed in the ileum upon *Rfx6* removal one week (Figure 4A-C, Table S3) and 3 months (Figure 4D-F, Table S4) after tamoxifen treatment by RNA sequencing. RT-qPCR were also performed in other segments of the small intestine and in the colon. (Suppl Figure 2). This strategy allowed us to identify early targets and later transcriptome changes. RNA-Seq revealed that 78 and 74 genes were respectively found up and down-regulated 1 week after tamoxifen treatment (Adjusted p-value ≤ 0.05) while these numbers increased to 1393 and 1483 after 3 months, suggesting adaptive responses. Overall, we found that transcripts for many intestinal hormones are almost undetectable (*Gcg, Pyy, Gip, Ghrl, Nts)* or decreased (Cck, *Sst, Nts, Sct*) in the small intestine of *Rfx6^ΔAdInt^* mice compared to controls (Figure 4B, E, Suppl Figure 2). *Gcg* and *Pyy* transcripts are decreased as well in the mutant colon (Suppl Figure 2B). In sharp contrast, *Tph1,* encoding the 5-HT/serotonin-generating enzyme in EC-cells, was up-regulated in the small and large intestine of *Rfx6^ΔAdInt^* mice. *Tph1* transcripts were increased as early as one week after *Rfx6* removal, in the duodenum (Suppl Figure 2A) and ileum (Figure 4B), and after one month in the jejunum and colon of *Rfx6^ΔAdInt^* mice (Suppl Figure 2B). Accordingly, *ChgA* expression, known to be enriched in EC-cells, was increased likewise upon *Rfx6* deletion (Figure 4E, Suppl. Figure 2B). In agreement with the transcriptomic data, we have not been able to find any SCT, GLP1-, GIP-, CCK-, or SST-expressing cell by immunofluorescence (data not shown). On the contrary, the number of serotonin (5-HT)-positive cells was increased already 1 week after *Rfx6* deletion compared to controls (Figure 5D). Consequently mucosal 5-HT content was higher in *Rfx6^ΔAdInt^* mice (Figure 5E). These observations suggest that the differentiation of endocrine cells producing peptides is impaired in absence of *Rfx6*, whilst the number of 5-HT cells is increased. Apart from the endocrine lineage, we did not observe any perturbation of the differentiation of other intestinal epithelial cells - including Paneth cells, goblet-cells or enterocytes - or of the crypt/villi morphogenesis (Suppl. Figure 3).

**Figure 4:**
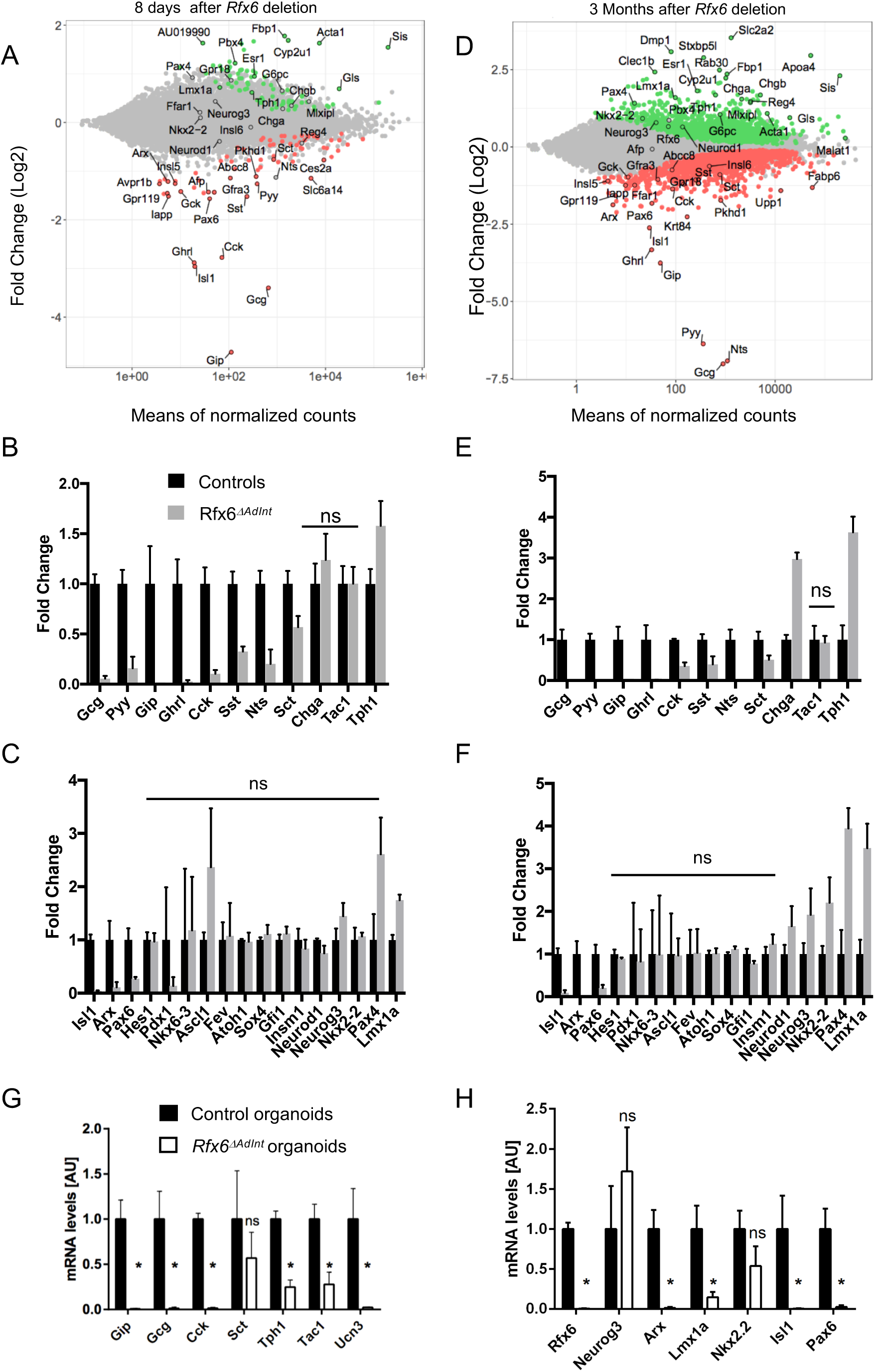
Gene expression profiling of the small intestine of adult Rfx6^ΔAdInt^ mice. **(A-F)** RNA-Seq analysis of the small intestine (ileum) of adult Rfx6^ΔAdInt^ mice 8 days (A-C) and 3 months (D-F) after Rfx6 deletion. n=3-4 animals were analyzed per group. (A, D) MA-plot representing the estimated log2 Fold-Change as a function of the mean of normalized counts. Green and red points represent significantly (adjusted pvalue ≤ 0.05) up- and down-regulated genes, respectively in Rfx6^Δ*AdInt*^ versus controls, 8 days (A) or 3 months (D) after Rfx6 deletion (RNA-Seq data). (B, C, E, F) Histograms showing RNA-Seq data for a selection of genes encoding enteroendocrine hormones or the rate limiting enzyme *Tph1* for serotonin synthesis (B, E) or transcription factors (C, F). All controls values are adjusted to 1. ns: non-significant, adjusted p-value >0.05 between controls and mutant. For all other genes adjusted p-value is ≤0.05. (G, H) RT-qPCR expression analysis of relevant genes in *Rfx6^ΔAdint^* duodenal organoids, 8 days after a 15 hours treatment with 1.25 µM Tamoxifen (4-OHT) to delete *Rfx6* gene. Data are represented as mean ± SD. Organoids were generated from n=4 mice per group. Statistics: non-parametric Mann*-*Whitney test, *** p≤0.001, ** p≤0.01, * p≤0.05.

**Figure 5:**
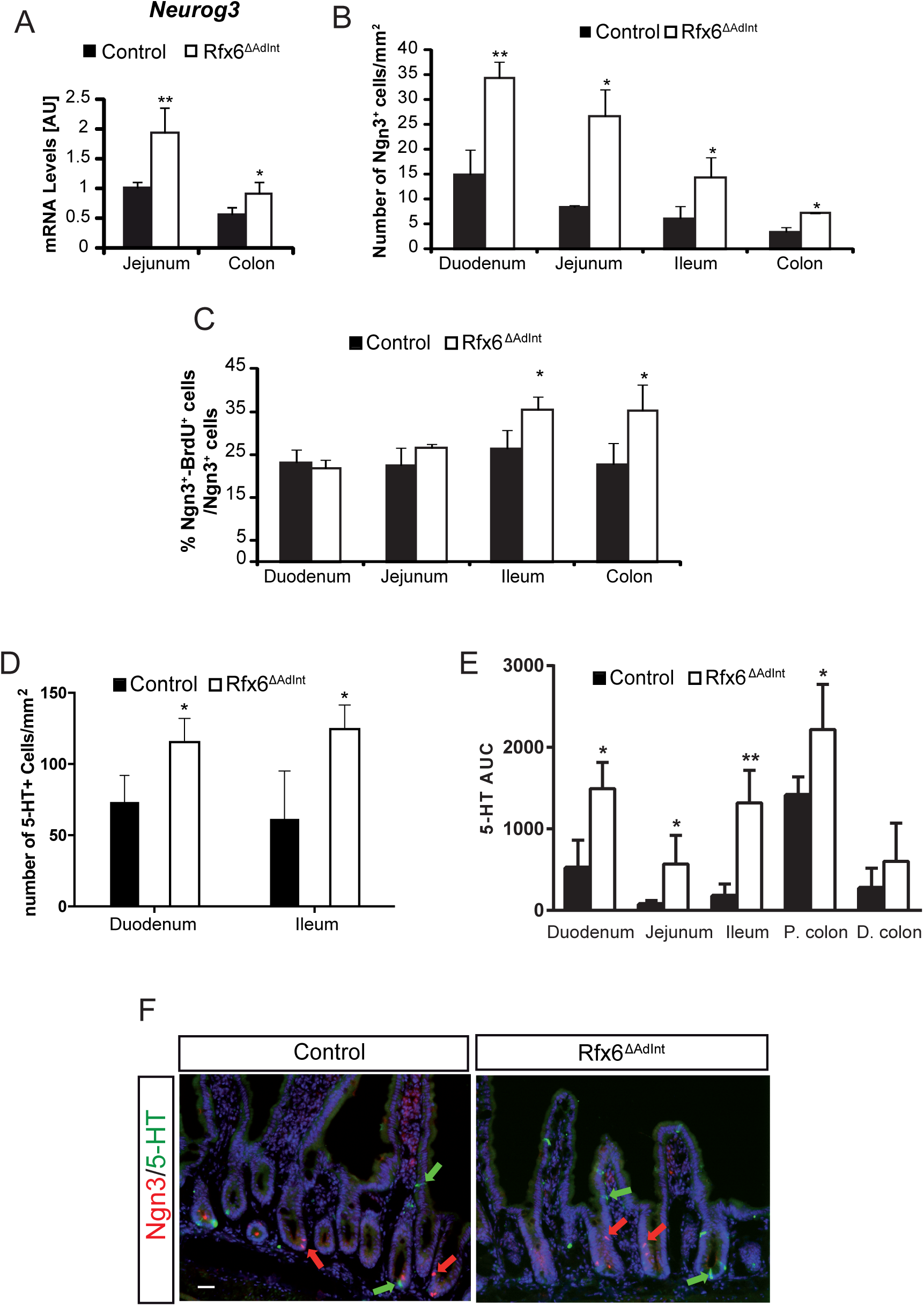
Increased number of Neurog3-positive endocrine progenitors and 5-HT-positive cells upon *Rfx6* deletion in the adult intestine. (A) RT-qPCR. Expression of *Neurog3* mRNA in the adult jejunum and colon three months after *Rfx6* deletion (n=4 mice per group). Unpaired t-test. (B) Quantification of the number of Neurog3-positive cells per mm^2^ of tissue in each segment of the adult intestine (n=3-4 mice per group) three months after *Rfx6* deletion. (C) Quantification of the number of proliferating Neurog3 cells expressed as the percentage of BrdU+/Neurog3+ cells on total Neurog3+ cells. (D) Quantification of the number of 5-HT-positive cells per mm^2^ of tissue in the duodenum and ileum of *Rfx*6^Δ*AdInt*^ and control mice, 8 days after *Rfx6* deletion (n=3 mice per group). (E) Mucosal 5-HT content analysis throughout the intestinal tract from day 11 of tamoxifen-treated control *Rfx*6^Δ*AdInt*^ mice (n =3-4 mice per group). Biopsies were sampled from Duodenum (Duo), Jejunum (Jej), Ileum, Prox. Colon (P. olon) and Dist. Colon (D. colon). 5-HT content were analysed on a HPLC-ECD and results stated as area under the curve (AUC). (F) Double immunofluorescence for Neurog3 and 5-HT on duodenal cryosections of control and Rfx*6^ΔAdInt^* mice (8 days post Tamoxifen treatment). Red and green arrows in D and E point to some Neurog3- and 5-HT-positive cells, respectively. Data are represented as mean ± SD; unpaired t-test was applied to calculate significance *p≤0.05; **p≤0.01. Scale bar represents 50 μm.

To determine the position of Rfx6 in the transcriptional network controlling enteroendocrine cell differentiation, we next examined the expression of transcription factors known to control enteroendocrine cell differentiation. We found that *Arx*, *Isl1* and *Pax6* are strongly decreased (Figure 4C, F, Suppl. Figure 2) as early as 1 week after tamoxifen treatment, in agreement with the impaired differentiation of peptidergic enteroendocrine cells and suggesting that Rfx6 acts upstream of these transcription factors. In contrast, the expression of several transcription factors was up-regulated (Figure 4C, F, Suppl. Figure 2) such as *Pax4* which is also up-regulated in *Rfx6*-deficent pancreatic beta cells. *Neurog3* transcripts are increased as well, suggesting that Rfx6 deletion perturbs endocrine progenitor cell development (Figure 5). Importantly, we also found that *Lmx1a* is increased upon *Rfx6* deletion in the adult intestine, and so it is the case for *Nkx2-2,* an upstream regulator of *Lmx1a,* 3 months after tamoxifen treatment. *Lmx1a* upregulation likely explains the increased expression of *Tph1*, the rate limiting enzyme for serotonin synthesis. RNA-Seq did not reveal any changes in expression for other endocrine transcription factors, such as *Insm1*, *Sox4*, *NeuroD1* or for *Atoh1* which is necessary for secretory lineage specification. Unaltered gene expression reflects either that the average gene expression levels are unchanged -which does not exclude variation in single cells- or that gene expression is independent of Rfx6. Taken together, our results suggest that Rfx6 controls transcriptional programs which promote the differentiation of enteroendocrine cells expressing peptide hormones (PE cells) while repressing the differentiation of enterochromaffin cells secreting serotonin (EC cells) in the digestive tract.

We next addressed the question whether Rfx6 similarly controls enteroendocrine cell differentiation in *Rfx6^ΔAdInt^* small intestinal organoids (Figure 4G, H). Briefly, tamoxifen was added to organoids cultures to remove Rfx6 and organoids were analyzed 8 days after treatment. As expected, RT-qPCR analysis confirmed the decreased expression of *Gip*, *Gcg*, *Cck* (*Sct* unchanged) upon *Rfx6* deletion. Unexpectedly, *Tph1* expression was also decreased, as *Ucn3* and *Tac1*, two other markers of EC cells (see last section results). In agreement with the decreased expression of EC enriched genes, *Lmx1a* was down-regulated as well in this culture system, suggesting that the consequences of Rfx6 removal on EC cells differentiation programs are different *in vivo* and *ex vivo*.

### 3.6 Enhanced expression of neoglucogenic and nutrient absorption machinery genes in the small intestine of mice upon Rfx6 deletion

To evaluate the long-term effect of impaired enteroendocrine cell function and lipid metabolism consecutively to Rfx6 deficiency, we examined the expression of intestinal genes that are not endocrine specific. Surprisingly we observed a strong and fast upregulation of genes involved in intestinal gluconeogenesis [46]. These include Fructose 1,6 biphosphatase (*Fbp1*) and Glucose 6 phosphatase (*G6pc*) as well as Glutaminase (*Gls*), the enzyme which hydrolyzes glutamine, the main gluconeogenic substrate in the small intestine (Figure 4 A, D, Table S4). The expression of the gene encoding the brush border enzyme sucrase-isomaltase (Sis), which hydrolyses dietary carbohydrates was also found to be up-regulated. A major long-term adaptation to intestinal Rfx6 deletion, not observed at 1 week, is the strong upregulation of the glucose transporter type 2 *Slc2a2* (also known as *Glut2*). Slc2a2 transporter is thought to facilitate the passage of dietary sugars in the blood stream [47] at the basolateral side of enterocytes. The expression of the sodium-coupled co-transporter *Sglt1,* which triggers glucose absorption at the apical membrane of enterocytes, was however not differentially expressed. We also found an up-regulation of *ApoA4*, an important chylomicron component, as well as of other apoliproteins such as Apoc3, Apoc2 and their upstream regulator, the transcription factor *Creb3l3/CREB-H* [48]. Together, our data suggest that *Rfx6^ΔAdInt^* mice adapt to overcome decreased food efficiency by promoting intestinal glucose and lipid absorption, as well as glucose production.

### 3.7 Increased number of enteroendocrine progenitors in intestinal crypts of *Rfx6^ΔAdInt^ mice*

Unexpectedly, when analyzing RNA-Seq data, we found increased levels of *Neurog3* transcripts in the ileum of adult *Rfx6^ΔAdInt^* mice 3 months after *Rfx6* deletion (Figure 4 C, F). Similarly, RT-qPCR data showed that *Neurog3* was up-regulated in the jejunum (2-fold) and in the colon (1.6-fold) compared to controls (Figure 5A) and in the duodenum as early as one week after *Rfx6* deletion (Suppl. Figure 2A). Similar results were observed in *Rfx6^ΔInt^* neonates (data not shown). Morphometric analysis revealed a 1.3-5 fold increase in the number of Neurog3-cells per mm^2^ in all segments of the small intestine as well as in the colon of *Rfx6^ΔAdInt^* mice (Figure 5B). To determine whether the increased pool of enteroendocrine progenitors resulted from an increased proliferation of endocrine progenitors, we quantified the percentage of Neurog3+/BrdU+ double-positive cells. Neurog3 cell proliferation was not affected in the duodenum and jejunum while was increased in the ileum and colon (Figure 5C). Neurog3-positive cells did not co-stain for 5-HT (Figure 5F) suggesting that the increased number of 5-HT-positive cells does not result from premature expression of 5-HT in enteroendocrine progenitors. Taken together, our results support that the increase of *Neurog3* transcripts in *Rfx6*-deficient intestine results from increased number of enteroendocrine progenitors.

### 3.8 Single cell transcriptomics revealed Rfx6 dependent genetic programs in the Peptidergic enteroendocrine and Enterochromaffin cell lineages

To gain knowledge into the mechanisms controlling enteroendocrine cell differentiation and diversity as well as into the role of Rfx6, we performed single-cell RNA sequencing of sorted EECs. Enteroendocrine cells were captured at various stages of their development, from endocrine progenitors to hormone expressing cells. Endocrine cells were purified from *Neurog3^eYFP/+^* mouse intestinal organoids. In this model, due to the stability of the eYFP protein, both crypt-based endocrine progenitors as well differentiating hormone-expressing EECs express the fluorescent reporter (Figures 6A and S4). After data filtering, 290 cells were analyzed. Shared Nearest Neighbor (SNN) clustering revealed eight groups of cells (Figures 6B and S5A) which were visualized in two dimensions using the Uniform Manifold Approximation and Projection (UMAP) algorithm. These clusters include 4 groups of progenitors (P1-4); P2-P4 expressing the endocrine progenitor marker *Neurog3* (Suppl. Figure 5A). Group P1 expresses markers of Paneth (*Defa 17, Lyz1*) and goblet (*Muc2, Tff3*) cells, suggesting multiple lineages priming in secretory progenitors as previously described [49]. The four other clusters represent subsets of Enterochomaffin cells (EC early and EC) expressing *Tph1* (Figure 6E and S5A), and Peptidergic Enteroendocrine cells (PE early and PE) expressing *Glc*, *Cck*, Sct, *Gip*, *Ghrl* and *Sst* (Figure 6F and S5A), at different stages of maturation. Clusters gene signature includes known and novel makers of EC and PE cells (Suppl. Figure 5A). Single-cell analysis confirmed that *Rfx6* is expressed both in endocrine progenitors as well as in more mature enteroendocrine cells of both EC and PE lineages (Figure 6D, O) and that single EECs can co-express several hormones genes (Figure 6O). Temporal resolution of transcriptome dynamics using diffusion map algorithm predicted cell state transition trajectories. Data analysis showed that endocrine progenitors differentiate in two branches: the EC and PE lineages (Figures 6C). Examples of genes expressed along the EC and PE branches are shown in Figure 6G-N and Suppl. Figure 6. Importantly, pseudotemporal ordering of single cells also revealed dynamics of transcription factors during EC or PE cell differentiation (Figures 6 and S6). Some transcription factors are clearly enriched in EC (*Lmx1a*, *Mnx1, Atf6, Glis3, Lhx1*) or PE (*Arx*, *Isl1*, *Pax6, Etv1*) lineage branches. Some other transcription factors (*Pax4*, *Neurod1*, *Nkx2-2*, *Insm1*, *Mlxipl),* like *Rfx6*, are found both in progenitors and more mature enteroendocrine cells (Suppl. Figure 6 Q-X) but are not specifically enriched in EC and PE lineages according to our selection criteria.

**Figure 6:**
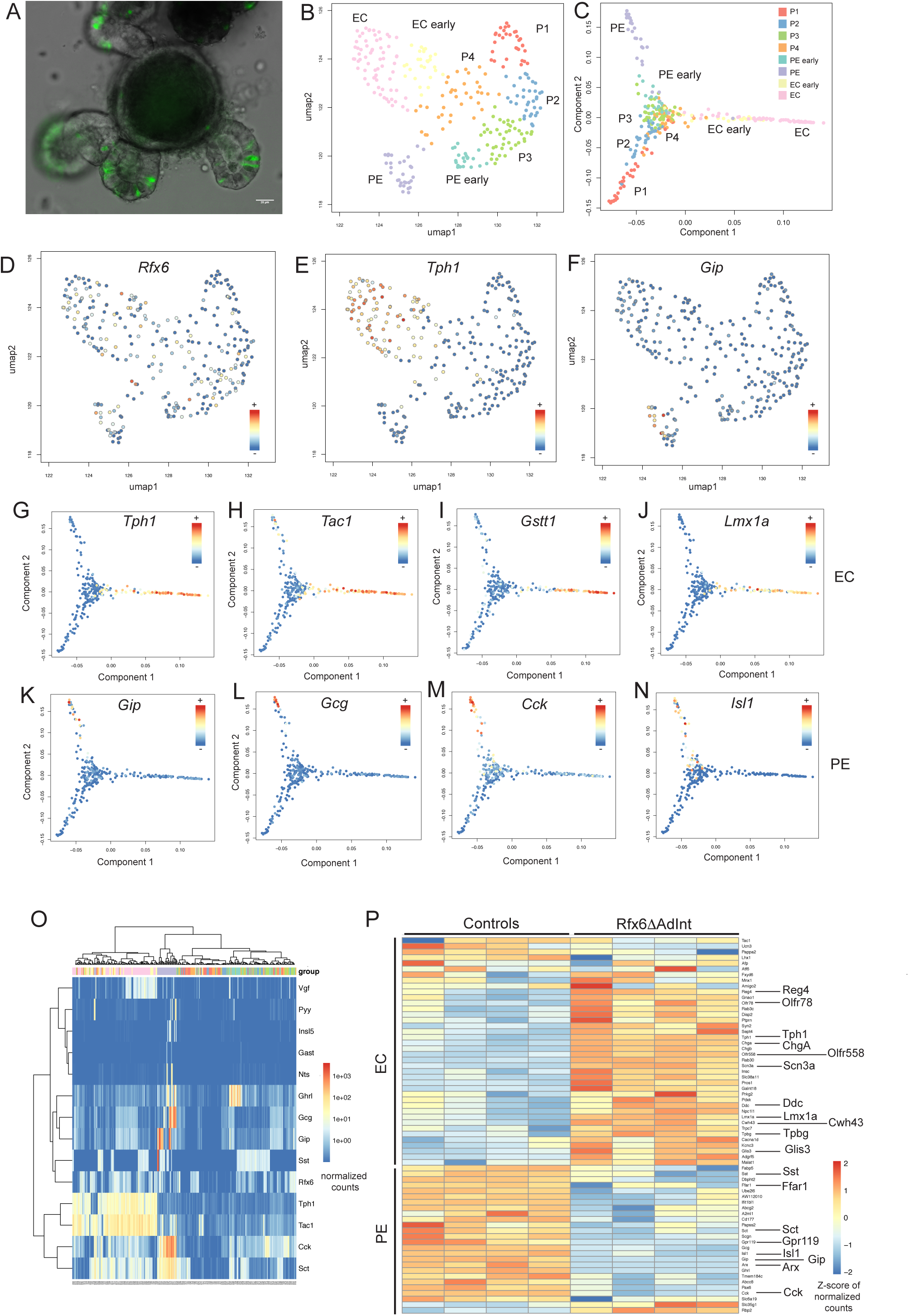
Single-cell RNA-Seq analysis of enteroendocrine cells of adult small intestine and Rfx6-dependent genetic programs. (A) Small intestinal organoid of Neurog3^eYFP/+^ mice (intrinsic eYFP fluorescence). eYFP labels Neurog3-positive endocrine progenitors and differentiated enteroendocrine cells. After organoid cell dissociation, fluorescence activated cell sorting (FACS) is used to isolate individual endocrine cells in 96 well plates. scRNA-Seq is then used to profile the cells. (B) Uniform Manifold Approximation and Projection (UMAP) representing the cell clusters. The 290 single cells were clustered based on the expression of 830 biologically variable genes. EC: Enterochromaffin cells; EC early: precursors of EC cells; PE: Peptidergic Enteroendocrine cells; PE early: precursors of PE cells; P1-4:groups of progenitors. (C) Cell clusters along the cell trajectory. Diffusion map, built from the log-expression of 830 biologically variables genes, was used to reconstruct the cell trajectory. (D-F) UMAP representing the expression of *Rfx6*, *Tph1* and *Gip* (normalized counts) in the 290 single cells. (G-N) Expression (normalized counts) of genes enriched in the EC (G-J) and PE (K-N) branches along the cell trajectory. (O) Heatmap (normalized counts) showing expression of hormone genes and Rfx6 in individual cells. (P) Heatmap (Z-score of normalized counts) showing expression of some markers of EC and PE lineages affected by Rfx6 deletion, in *Rfx6*^Δ*AdInt*^ (3 months after Rfx6 removal) and control conditions. Shown is the expression (row-wise Z-score), in *Rfx6*^Δ*AdInt*^ and control conditions, of markers of EC populations (EC and/or EC-early populations) and PE populations (PE and/or PE-early populations) which are differentially expressed after Rfx6 deletion.

Our data show that Rfx6 differentially controls genetic programs regulating PE and EC cells, which appeared to be the two main EEC lineages. We thus determined, among the genes defining the signature of PE and EC lineages, which one are actually regulated by Rfx6 in the adult small intestine (Figures 6P ad Supll. Figure 5B). As expected, most PE enriched genes, like hormone genes (e.g *Sct*, *Gcg*, *Gip*, *Sst*, *Cck*), related transcription factors (e.g. *Arx*, *Isl1*, *Pax6*), GPCRs (e.g *Gpr119*, *Ffar1*) as well as other PE markers (*Abcc8*, *Scgn*, *Fapb5*) are down regulated in absence of Rfx6. Surprisingly rare PE enriched genes are either up-regulated (*Rbp2*) or have their expression unchanged (*Gpr112*, *Rbp4*, *Scg3*, data not shown). Furthermore, the vast majority of EC enriched genes are up-regulated in absence of Rfx6, including endocrine markers (*Tph1*, *ChgA*, *Scn3a*), transcription factors (*Lmx1a*, *Glis3)*, GPCRs (*Olfr558*, *Olfr78)* or other markers (e.g *Cacna1d*, *Disp2*, *Cwh43*). However, the expression of highly specific markers of the EC lineage was not affected (e.g *Tac1*, *Ucn3*, *Lhx1*, *Mnx1*, *Atf6*) or decreased (*Pappa2*, *Ly6a*) by the loss of Rfx6. Taken together, these results suggest that Rfx6 regulates specific genetic programs in the EC and PE lineages, while others genes are independent of Rfx6.

## 4. DISCUSSION

Mutations in *RFX6* are causal of Mitchell-Riley syndrome in human. This syndrome is characterized by neonatal diabetes and intestinal failures including malabsorption [21]. Previous studies showed that Rfx6 controls beta-cell development in the mouse embryo as well as insulin secretion in adult mouse and human beta cells [21, 22, 31]. However, the function of Rfx6 in the mouse intestine was unknown. In this study we show that, as in the pancreas, Rfx6 is expressed in Neurog3-endocrine progenitors and persists in all hormone-expressing cells in the intestine, and not only in Gip-expressing cells as previously thought [23]. Analysis of mice with a constitutive or inducible intestinal deletion of Rfx6 resulted, in all cases, into a severe loss of enteroendocrine cells expressing peptide hormones while enterochromaffin cells were still present and genes controlling serotonin-production were increased. Unlike newborns that die, adult mice can cope with the induction of this massive reduction of peptidergic EECs. However, intestinal absorption is perturbed leading to impaired food efficiency. RNA-Seq revealed enhanced expression of neoglucogenic and nutrient absorption machinery genes that we hypothesize to be a compensatory mechanism to cope with the malabsorption. Importantly, using single-cell transcriptomics we found that EECs differentiate from endocrine progenitors according to two cell trajectories: the Enterochromaffin cells (EC) and Peptidergic Enteroendocrine (PE) cells. Similar conclusions were drawn recently in a single-cell study [50]. Identification of specific gene signatures and dynamics in the EC and PE lineages allowed us to determine their respective dependence on Rfx6. We propose that Rfx6 positively regulates genetics programs leading to PE cells differentiation, and represses 5-HT production, as well as other, but not all, EC cell specific programs (Figure 7).

**Figure 7:**
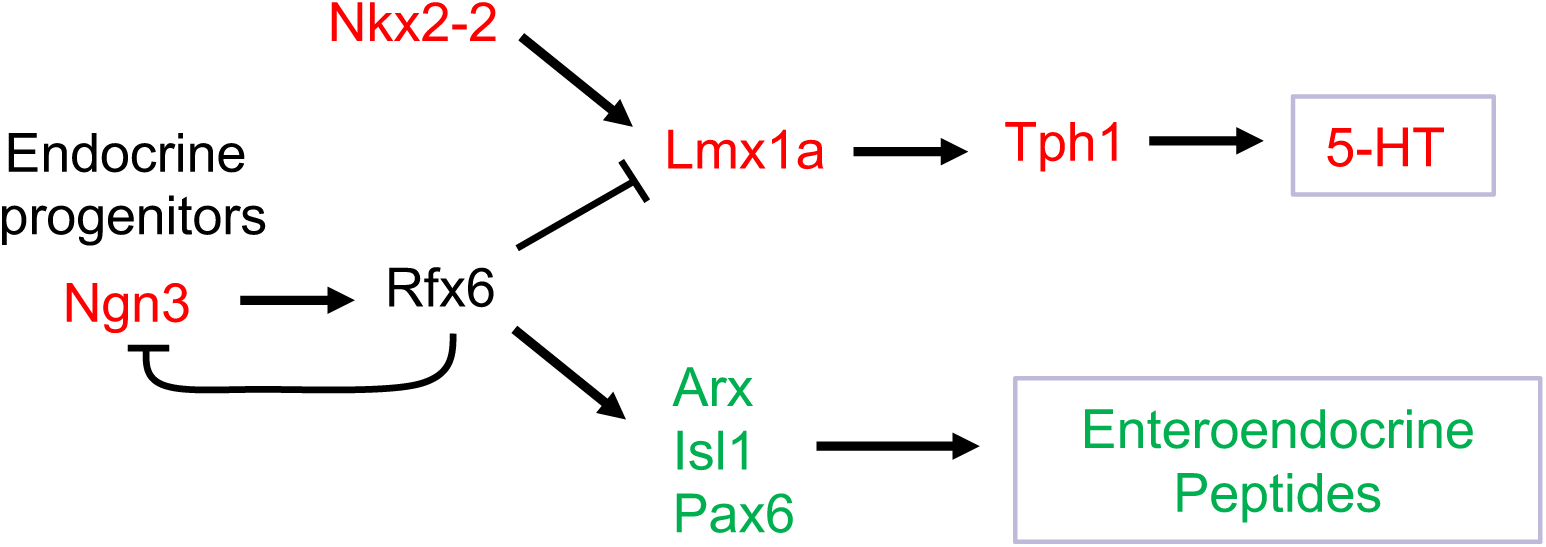
Proposed model for the role of Rfx6 in enteroendocrine lineages specification in the adult intestine. Neurog3 triggers the endocrine fate. *Rfx6* is then activated in endocrine progenitors downstream of Neurog3 and represses *Neurog3* transcription allowing differentiation to proceed. Rfx6 promotes peptidergic endocrine (PE) cell differentiation upstream of *Arx*, *Isl1* and *Pax6*. Rfx6 represses, upstream of *Lmx1a*, a genetic program leading to *Tph1* expression and 5-HT production. Green and Red: genes respectively down or up-regulated in the *Rfx6^ΔAdInt^* compared to wild-type controls.

In addition to the severe alteration in Peptidergic Enteroendocrine cell differentiation, our studies demonstrated that constitutive inactivation of Rfx6 leads to gastric heterotopia in the embryonic mouse intestine. Heterotopic gastric mucosa was reported in only 3 patients carrying compound heterozygote mutations in *RFX6* [29, 30]. These patches of gastric heterotopia were revealed after histological analysis of intestinal resection to treat hemorrhagic ulcers. It has been proposed that mucosa alteration could be caused by acid secretion of the ectopic gastric tissue [30]. It is likely that gastric heterotopia is a general feature of patients with Rfx6 mutation, since anemia and melena were reported in many cases and could result from intestinal bleeding [30]. In one case, ectopic pancreatic tissue was also reported in the jejunum [30], a feature that we did not observe in *Rfx6*-deficient mice. Importantly, we observed gastric tissue only in the intestine of mice with a constitutive deletion of *Rfx6*, never when *Rfx6* was deleted in the endocrine lineage or when Rfx6 was inactivated in the adult intestine. These observations suggest that gastric heterotopia is unrelated to the impairment of the endocrine lineage but rather results from an early function of Rfx6 in mouse endoderm patterning, similarly to what has been suggested in Xenopus [24]. Nevertheless, we did not observe any gross defect of intestinal or gastric organogenesis, suggesting functional redundancy. The patchy pattern of gastric tissue in *Rfx6*-deficient intestine might thus be explained by the lack of functional compensation in some pluripotent endodermal stem cells that consequently lost or did not acquire the proper regional identity.

*Rfx6* inactivation led to an increased expression of *Tph1*, which is critical for 5-HT biosynthesis in enterochromaffin cells, at all stages and conditions studied *in vivo* throughout the intestine. Furthermore, in the adult intestine, we measured an increased number of 5-HT-positive cells and mucosal 5-HT content in the small intestine upon Rfx6 removal. We thus propose that Rfx6 represses 5-HT biosynthesis. Of note, we found that the expression of *Lmx1a,* shown previously to control, downstream of *Nkx2-2,* the expression of *Tph1* [51], was up-regulated. This suggests that increased *Tph1* results from increased *Lmx1a*. By contrast, *Nkx2-2* transcripts were unaffected one week after Rfx6 deletion, suggesting that Rfx6 does thus not repress *Lmx1a* via Nkx2-2. However, since 3 months after the tamoxifen treatment, *Nkx2-2* expression was increased, we cannot exclude a contribution of Nkx2-2 in *Lmx1a* and *Tph1* up-regulation at a later stage. Importantly, the unchanged expression of several specific markers of the EC lineage, identified in our single cell transcriptome study of enteroendocrine cells, such as *Ucn3*, *Tac1*, *Atf6*, *Lmx1a* and *Mnx1,* suggests that Rfx6 does not act as a genetic switch promoting PE and inhibiting EC destiny. Rather, Rfx6 represses the genetic program leading to 5-HT biosynthesis and possibly others specific features of EC cells. However, as some of these genes are also expressed outside of enterochromaffin cells, such as *Ucn3* or *Tac1* which are found in enteric neurons [52], it is possible that we could not detect variation in their mucosal expression when performing bulk RNA-Seq on the entire intestinal tissue. Of note, the cellular origin of the increased expression of *Lmx1a* and *Tph1* remains unclear. One hypothesis is that the loss of Rfx6 results into the ectopic induction of the serotonergic program in the PE lineage. Another possibility would be that Rfx6 modulates the levels of 5-HT in EC cells by controlling the expression of *Lmx1a* and *Tph1*. The absence of cells co-expressing Neurog3 and 5-HT in *Rfx6^ΔAdInt^* mice suggests that Rfx6 does not represses the serotonergic program in enteroendocrine progenitors but rather at later stages of EEC cell differentiation.

Our study also sheds light on the position of Rfx6 in the regulatory network of transcription factors controlling EECs differentiation. We found that *Pax6*, *Isl1*, and *Arx* are down-regulated, while *Pax4* is up-regulated by the inactivation of Rfx6 suggesting that Rfx6 operates upstream of these transcription factors. In *Arx*-deficient mice, the expression of *Gcg*, *Gip*, *Cck* and *Nts* was reduced [14, 15], suggesting that Rfx6 regulates the expression of these hormone-encoding genes through the regulation of *Arx*. Pax4 and Arx are known to repress each other in the developing pancreas [53]. However, this does not seem to be the case in the gut. Indeed, *Pax4* expression is unchanged in *Arx*-deficient intestine. Thus, increased *Pax4* expression, in absence of Rfx6, is probably not the consequence of *Arx* down-regulation. Nevertheless, *Pax4* was shown to be required for EC cell differentiation [14] in agreement with the maintenance of EC cells in Rfx6-deficient mice. Rfx6 is thus turned on in endocrine progenitors and operates downstream of Neurog3 to regulate a network of transcription factors including Lmx1a, Pax4, Arx, Pax6 and Isl1 which will control later steps of EEC differentiation. Our study revealed that *Neurog3* expression and the number of Neurog3-positive endocrine progenitor cells increased upon induction of Rfx6 deletion in both the small intestine and colon. One explanation for this increase would be that Rfx6 normally represses Neurog3 expression in late endocrine progenitors. Alternatively, endocrine progenitors accumulate because they cannot differentiate further into Peptidergic enteroendocrine cells. Unexpectedly, when we studied the consequences of the induction of Rfx6 deletion in small intestine organoid cultures, we found that development of all EECs was impaired including EC cells. In line with this observation, the expression of *Lmx1a* and *Tph1* was down-regulated *ex vivo* upon Rfx6 deletion. A similar conclusion was reported recently in a study where Rfx6 was inactivated directly, *ex vivo*, in mouse intestinal organoids [50]. We propose that these *in vivo/ex vivo* differences on the role of Rfx6 in EC cell development could reflect limitations of the organoid culture system to faithfully mimic alternative cell differentiation pathways when genetic networks are perturbed. We cannot exclude that the promotion of serotonergic programs in the absence of Rfx6 results from an adaptation due to a systemic effect or an effect of the microbiota that would not occur *ex vivo*. However, we do not favor this hypothesis as we saw the induction of *Lmx1a* and or *Tph1* when Rfx6 is lacking, not only in the adult but also in the embryonic intestine and in the developing stomach suggesting a general function of Rfx6 in repressing enterochromaffin cell differentiation.

In conclusion, this study revealed that Rfx6 is essential, downstream of Neurog3, for the differentiation of peptidergic enteroendocrine cells during development and in the adult. Rfx6-dependent impairment of enteroendocrine cells perturbs food efficiency. In addition, we show that Rfx6 also represses genetic programs leading to the biosynthesis of 5-HT. These findings shed new light on the molecular mechanisms underlying intestinal malabsorption and energy metabolism in RFX6 deficiency in humans

## AUTHOR CONTRIBUTION

J.P. designed and performed most of the mouse experiments, analyzed the data and wrote the paper, C.V. performed bioinformatic analysis, interpreted results and wrote the paper. F.B. and A.D.A. performed single cell experiments, interpreted results and wrote the paper. F.B., A.D.A. and N.P performed all organoids experiments and interpreted results. S.G., L.N., M.L.L. M.E. and C.T.C helped initially with the design and realization of single cell experiments. C.K., S.J.C. and N.M. contributed to the bioinformatic analysis. M.M.L performed serotonin measurements. J.P. and A.D.A. performed morphometric analysis. A.M., A.B., P.S. and V.S., performed experiments and contributed to data analysis. C.L assisted with the immunostaining experiments. T.W.S provided conceptual input. G.G. oversaw the entire project, designed experiments, analyzed data, wrote the paper and obtained financial support.

## ACKNOWLEDGEMENTS

The authors thank the members of the Genomeast platform for RNASeq and Doulaye Dembele for initial analyses of the data, of the Cell sorting facility, of the Mouse Clinical Institute Metabolic Exploration Plateform (MCI, Strasbourg, France) for their help with the metabolic analyses. We thank N. Messaddec (IGBMC), for Electron microscopy and Martine Poulet for technical assistance and Tao Ye (IGBMC) for submission of data to GEO. We thank Miriam Ejarque (IGBMC), Catherine Tomasetto (IGBMC), Natalie Terry (Division of GI, Hepatology, and Nutrition Children’s Hospital of Philadelphia) and Klaus Kaestner for helpful discussions. This study was funded by the Novo Nordisk Foundation (Challenge Grant NNF14OC0013655) and by Grants from the Agence Nationale pour la Recherche (ANR Rfx-PancInt) and from the Foundation pour la Recherche Médicale (FRM). IGBMC is supported by the grant ANR-10-LABX-0030-INRT, a French State fund managed by the Agence Nationale de la Recherche under the frame program Investissements d’Avenir ANR-10-IDEX-0002-02.

---

**Figure S1:**
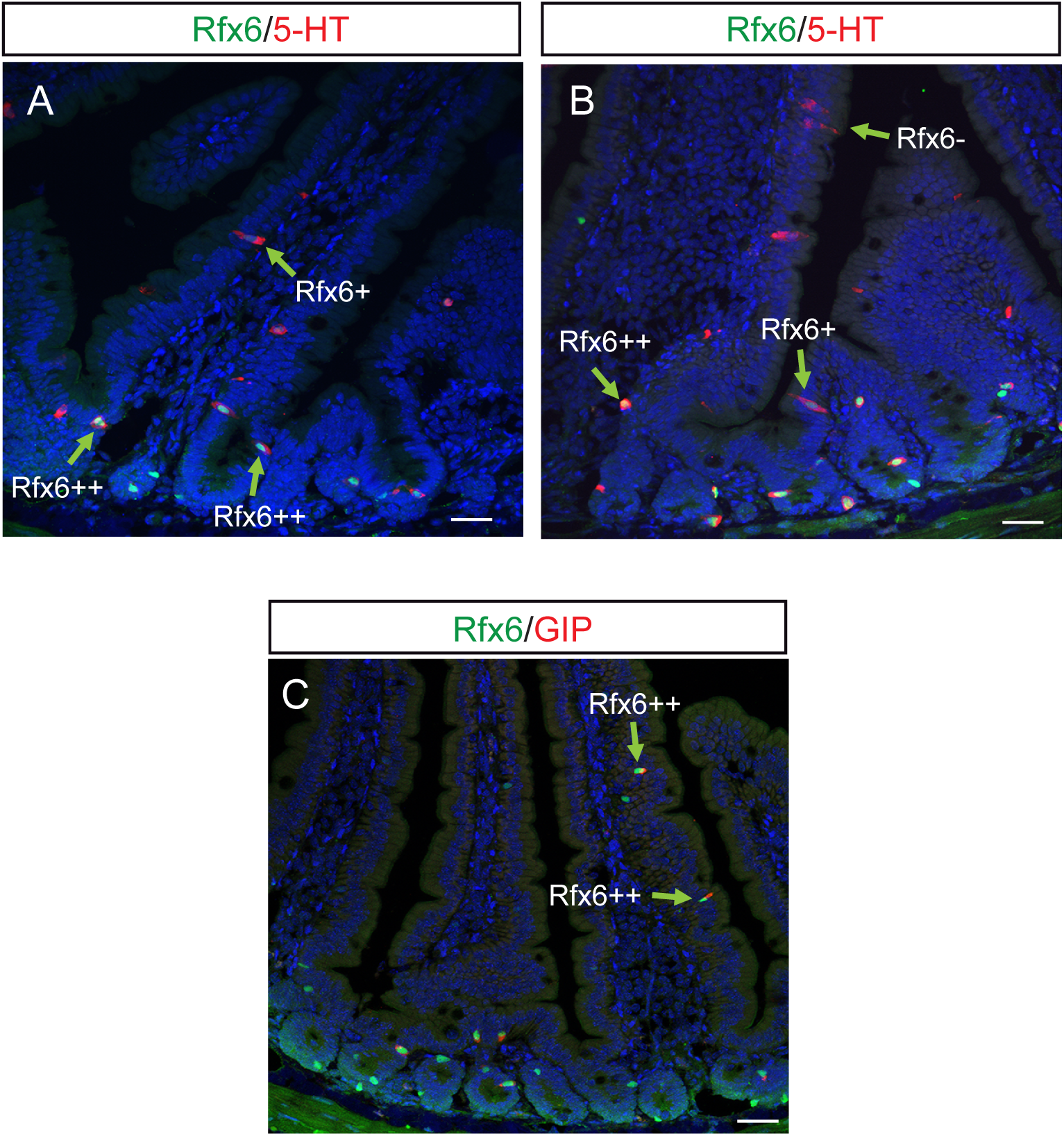
Expression of Rfx6, 5-HT and GIP in adult mouse duodenum. (A-B) Immunofluorescence staining for Rfx6 (green) and 5-HT (red). (C) Immunofluorescence staining for Rfx6 (green) and GIP (red). Arrows point to enteroendocrine cells expressing Rfx6 at different levels. Scale bars:50 µM.

**Figure S2:**
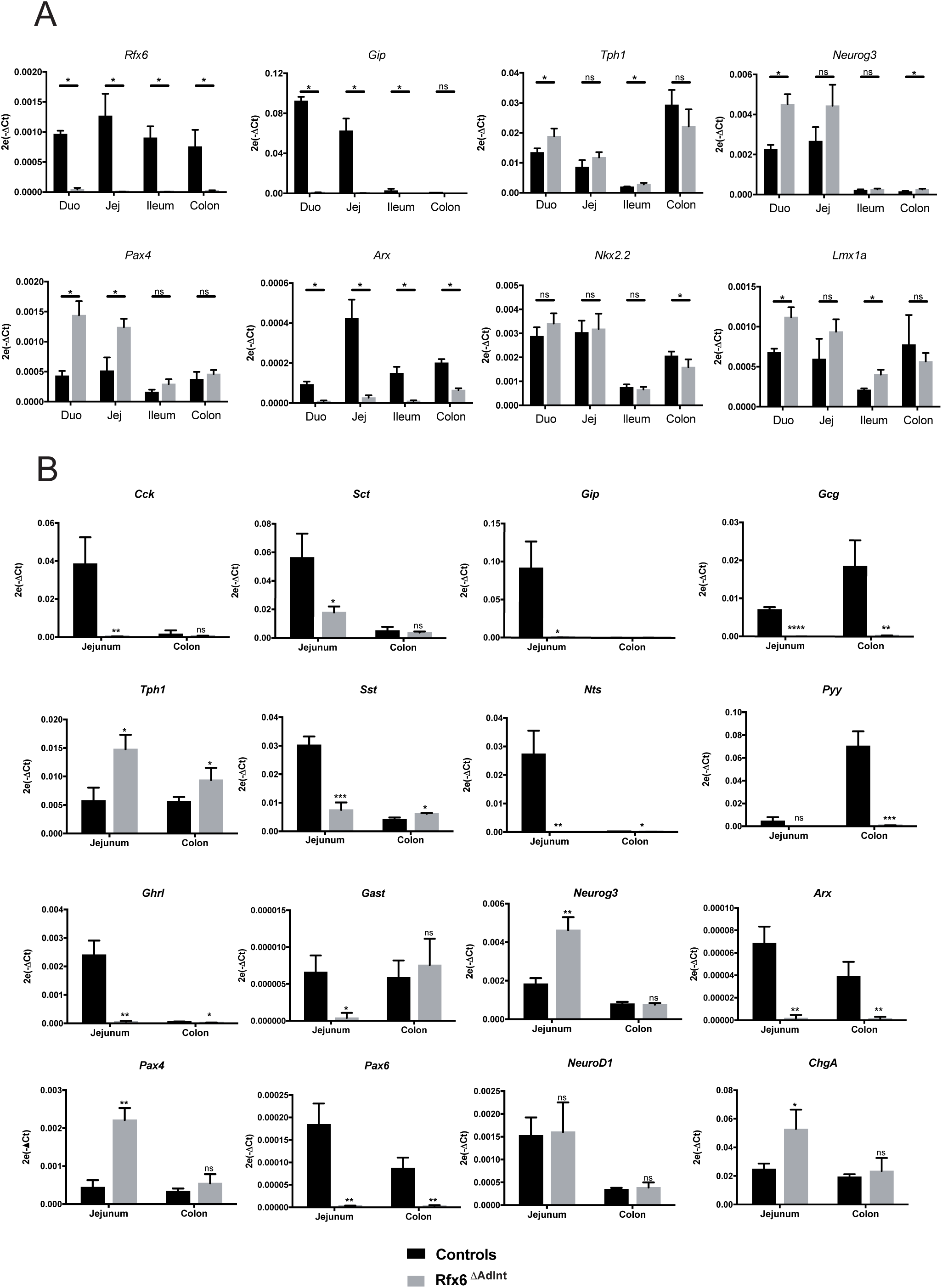
Loss of Rfx6 in the adult intestine strongly impairs enteroendocrine cell differentiation except serotonin expressing cells. RT-qPCR experiments to determine the expression of selected genes in controls (black) and Rfx6^ΔAdInt^ (grey) mice in the small and large intestine. Analysis were performed 8 days (A, n=4 animals per group) or one month (B, n=3 animals per group) after Rfx6 deletion. Statistics, Mann Whitney (A) or unpaired t-test (B). Relative mRNA levels are represented as 2^-ΔCt^ means ± SD; *** p≤0.001, ** p≤0.01, * p≤0.05, ns: not significant.

**Figure S3:**
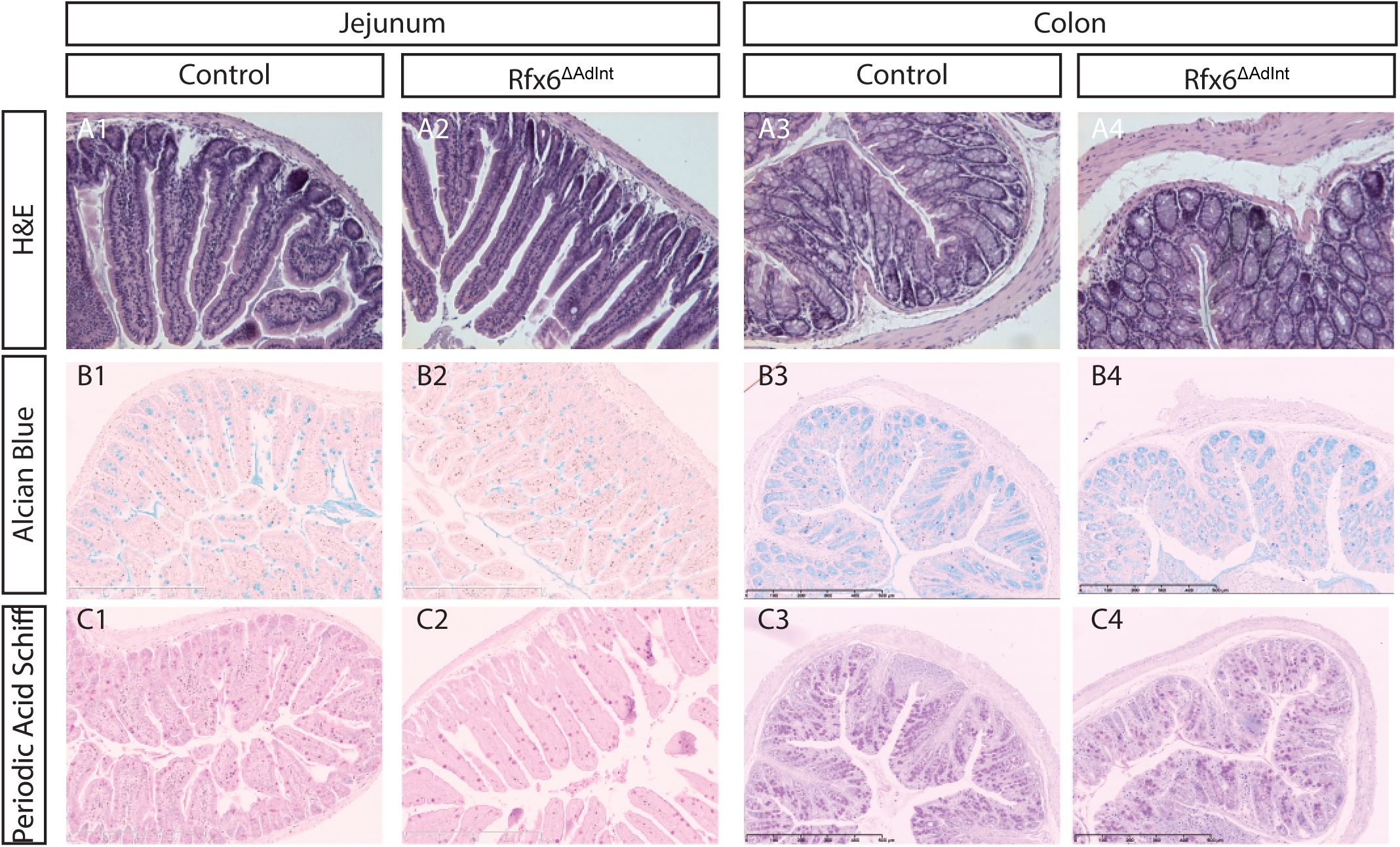
Rfx6 deletion in the adult intestine does not lead to any perturbation of the crypt/villus morphogenesis, differentiation of Paneth, goblet cells or enterocytes. Set of pictures representing the jejunum (columns 1-2) and colon (columns 3-4) of controls (columns 1&3) and Rfx6^ΔAdInt^ (columns 2&4). (Row A) Haematoxylin and Eosine staining (HE). (Row B) Alcian blue staining of acidic mucins. (Row C) Periodic Acid Schiff staining of mucins.

**Figure S4:**
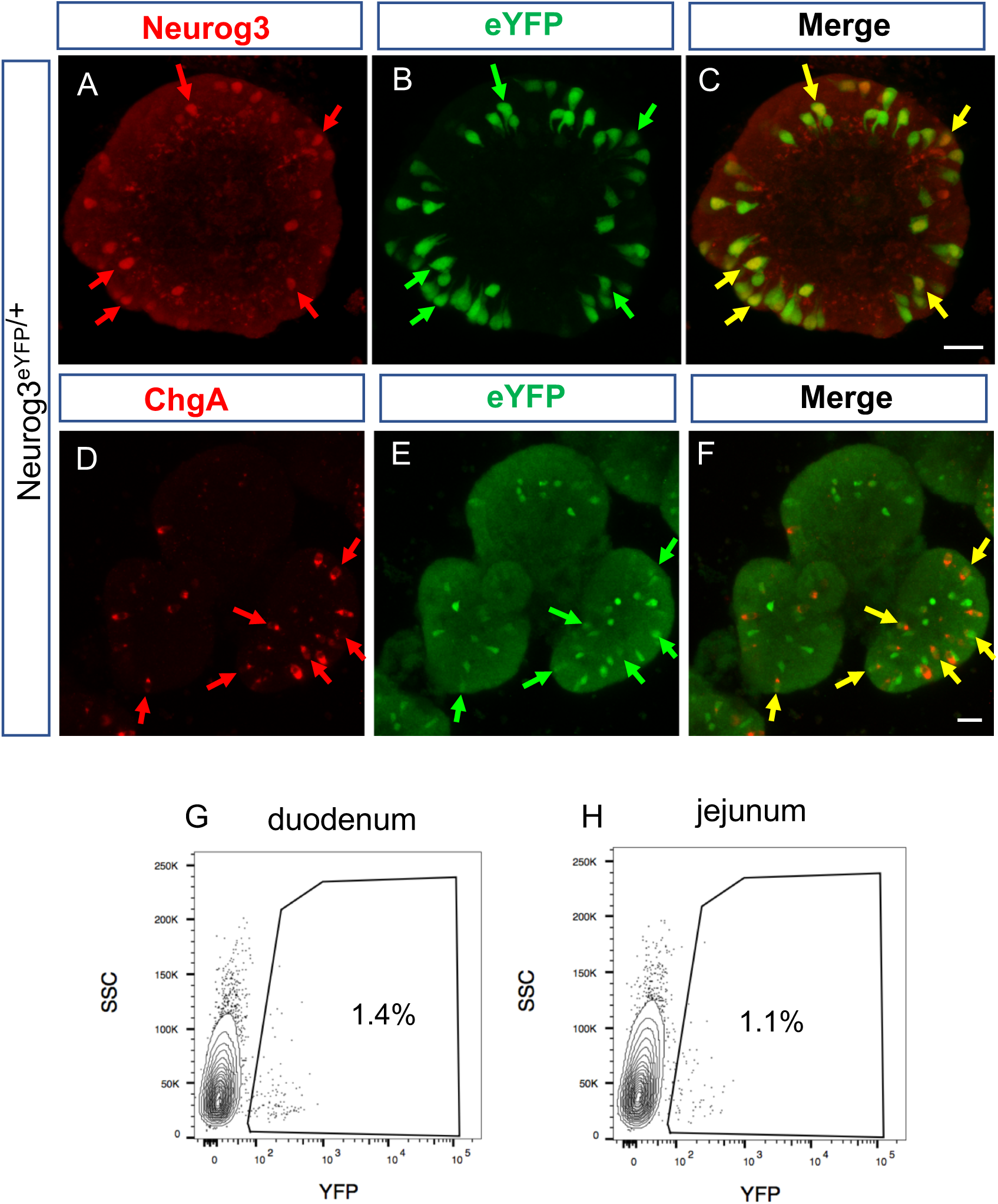
Histological and FACS analysis of endocrine cells in small intestine organoids of Neurog3^eYFP/+^ mice. (A-G) Fluorescent confocal microscope pictures of small intestine organoids co-labeled (whole-mount) with eYFP (Green) and Neurog3 or ChgA (Red), depicting expression of eYFP reporter in endocrine progenitor cells (Neurog3-positive) and maintenance in their differentiated progeny (ChgA-positive). Yellow arrows point to co-labeled cells. Scale bar: 25 µM. (H-I) Typical FACS profile depicting purification of eYFP+ cells from duodenal (H) and jejunal (I) Neurog3^eYFP/+^ organoids. SSC: (Side Scater).

**Figure S5:**
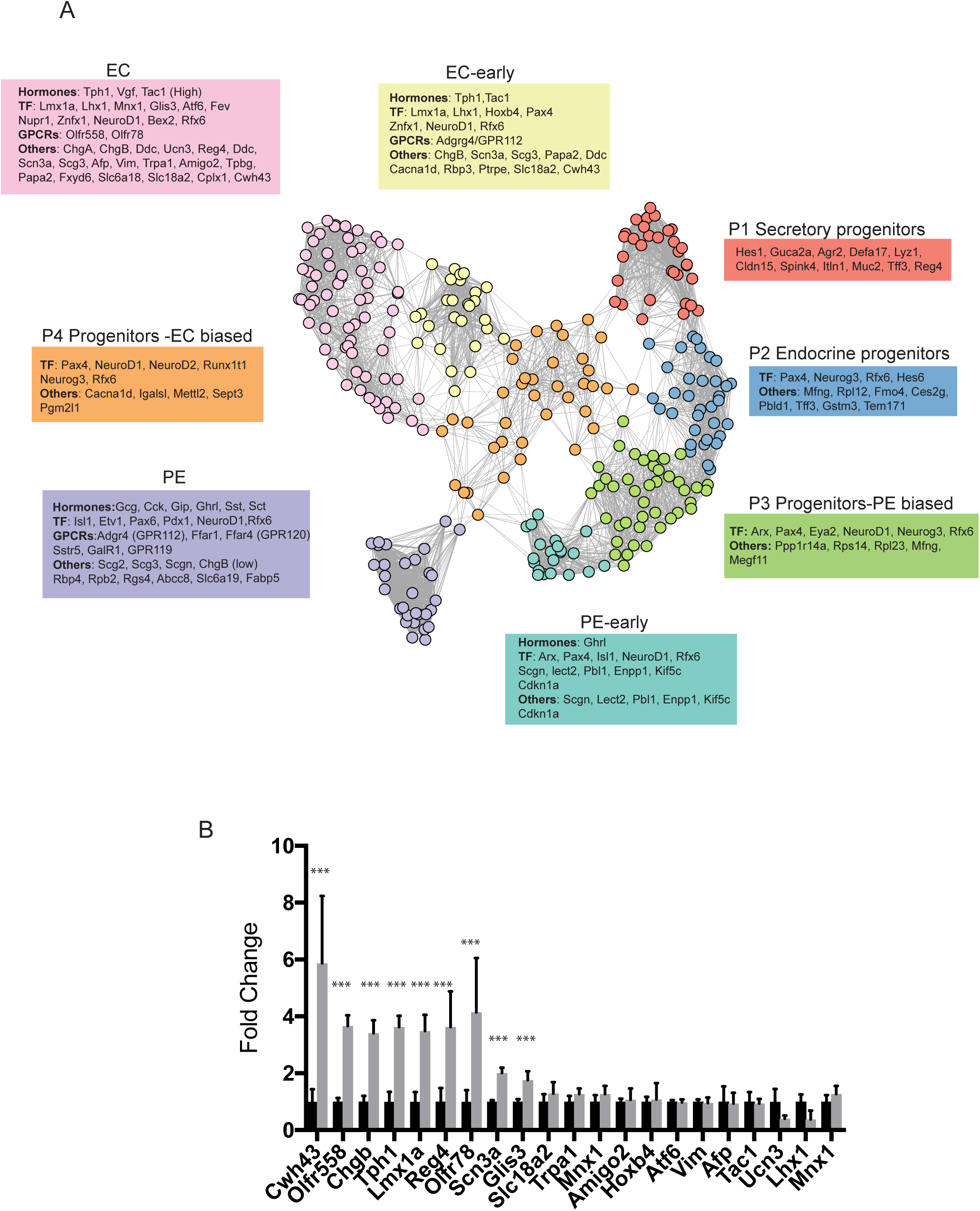
(A) Uniform Manifold Approximation and Projection (UMAP) representing the 8 cell clusters and their markers. The 290 single enteroendocrine cells (eYFP+ sorted from Neurog3 - eYFP small intestine organoids) were clustered based on the expression of 830 biologically variable genes. EC: Enterochromaffin cells; PE:Peptidergic enteroendocrine cells; P1-P4: Progenitor cell populations. **(B) Increased expression of several EC genes upon Rfx6 deletion**. RNA-Seq analysis of adult ileum of Rfx6^ΔAdint^ (grey boxes) and control (black boxes) mice, 3 months after tamoxifen treatment. Data shown are a selection of genes found strongly enriched in the EC branch in single cell transcriptomic of wild-type enteroendocrine cells. n=3-4 animals were analyzed per group; ***; adjusted p-value ≤0.001.

**Figure S6:**
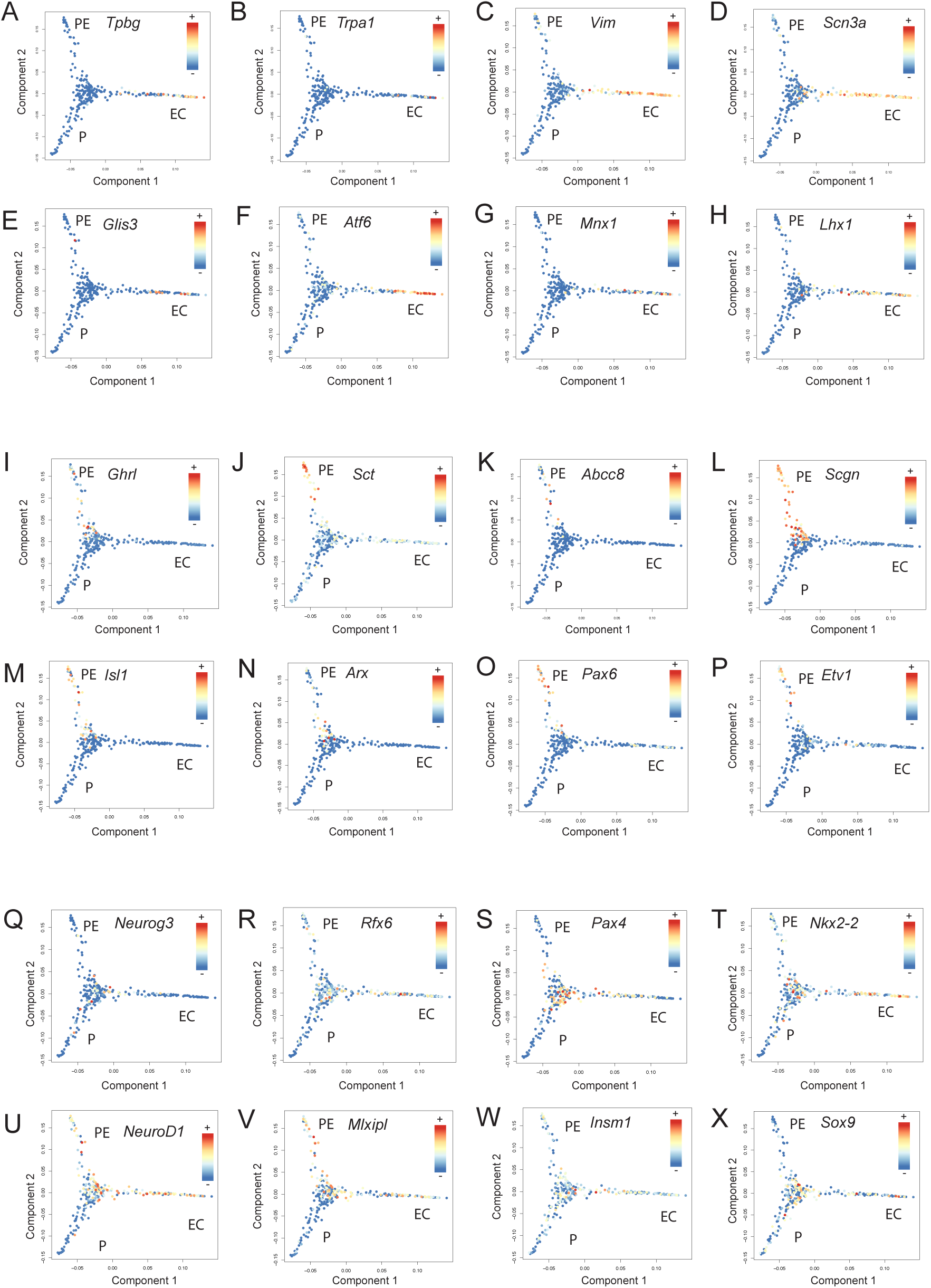
Expression of selected genes along cell trajectories during enteroendocrine cells differentiation from progenitors to enterochromaffin and peptidergic enteroendocrine branches. Diffusion map, built from the log-expression of 830 biologically variables genes, was used to reconstruct the cell trajectory. Expression (normalized counts) of some genes enriched in the enterochromaffin branch (A-H) in the peptidergic branch (I-P) and of a selection of transcription factors (Q-X). P: progenitors; EC: enterochromaffin cells, PE:peptidergic enteroendocrine cells.

**Table S1 :**
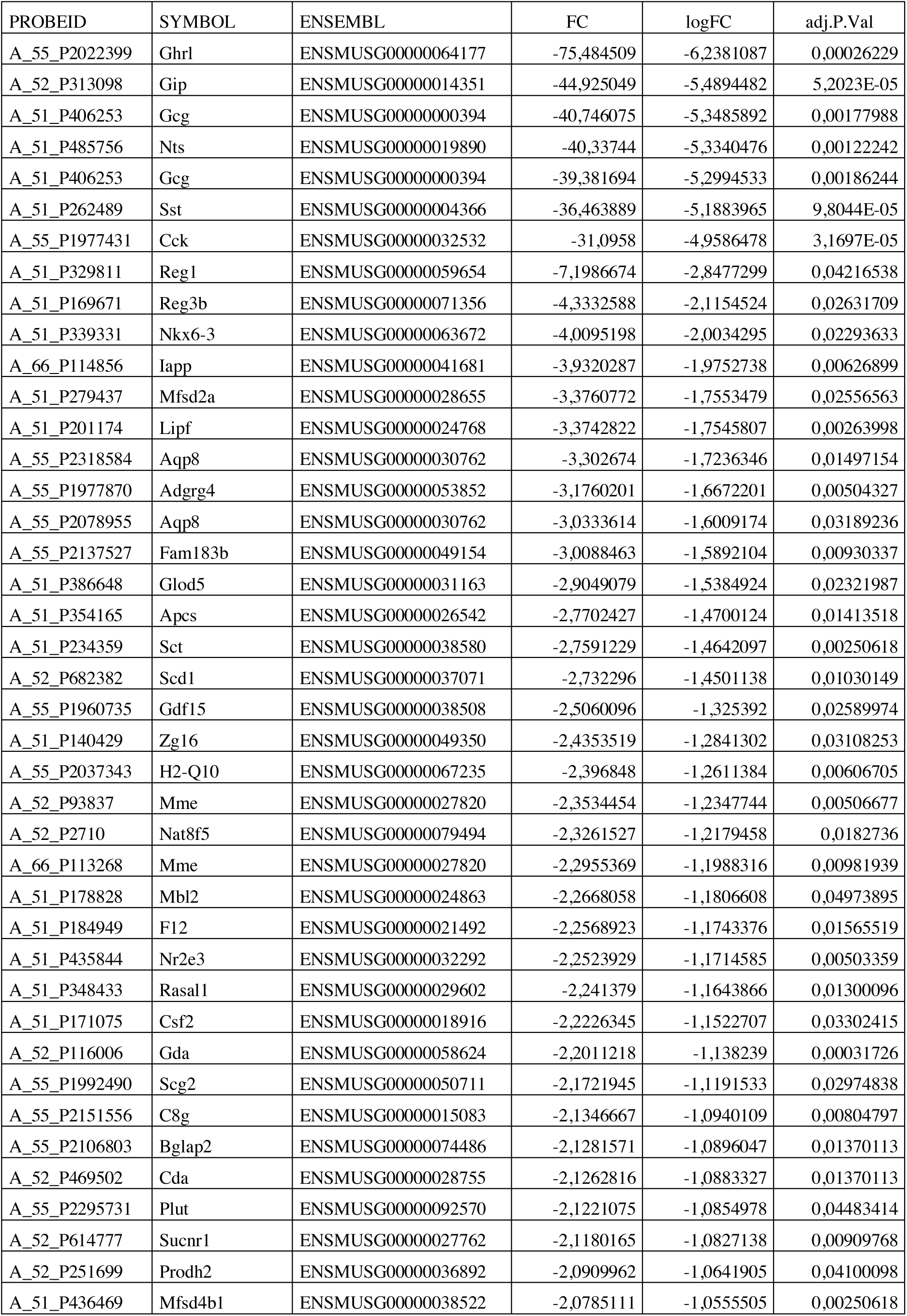
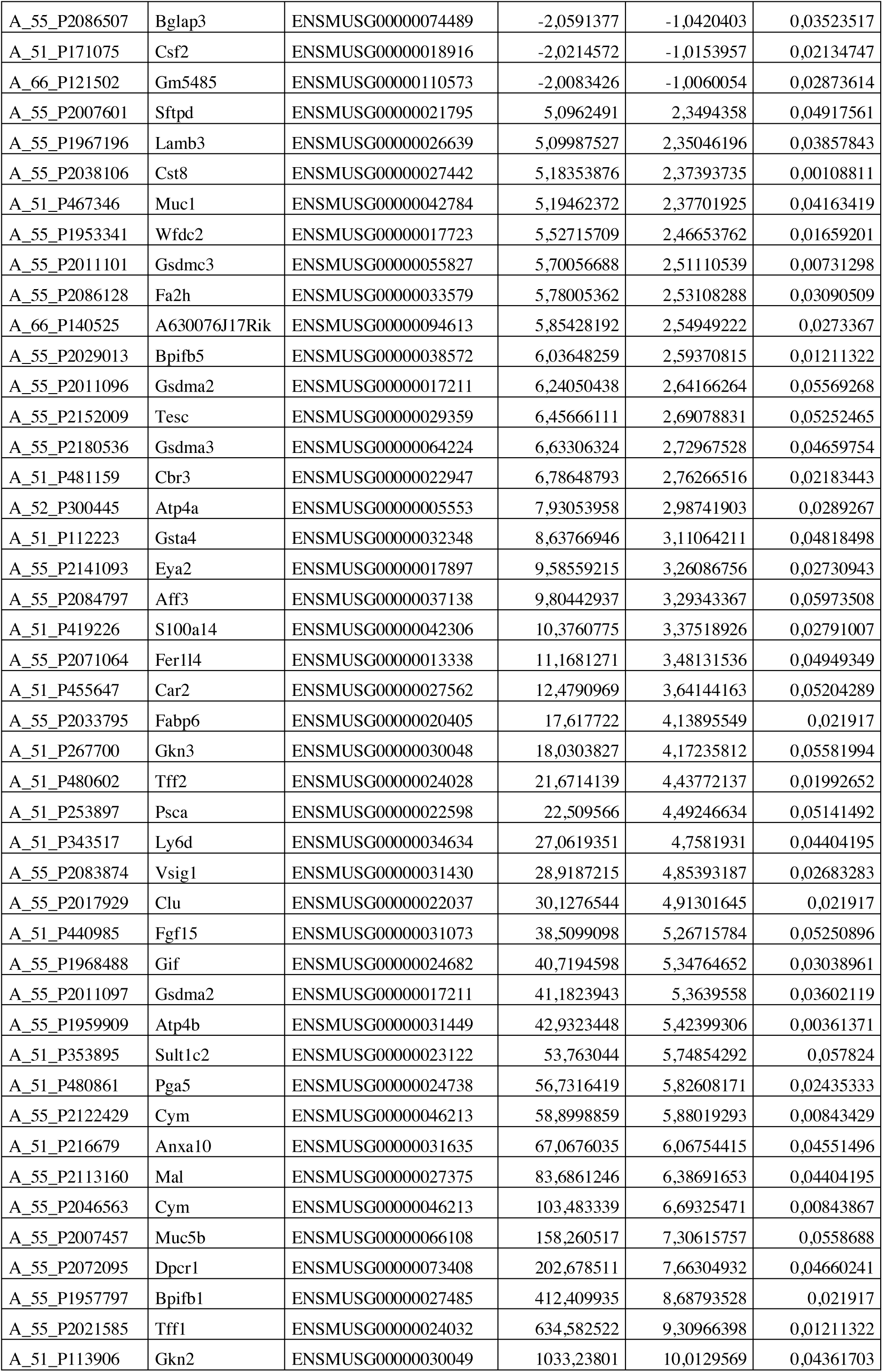
List of genes differentially regulated in Rfx6^-/-^ small intestines (E18.5) compared to controls ordered by Fold Change.

**Table S2 :**
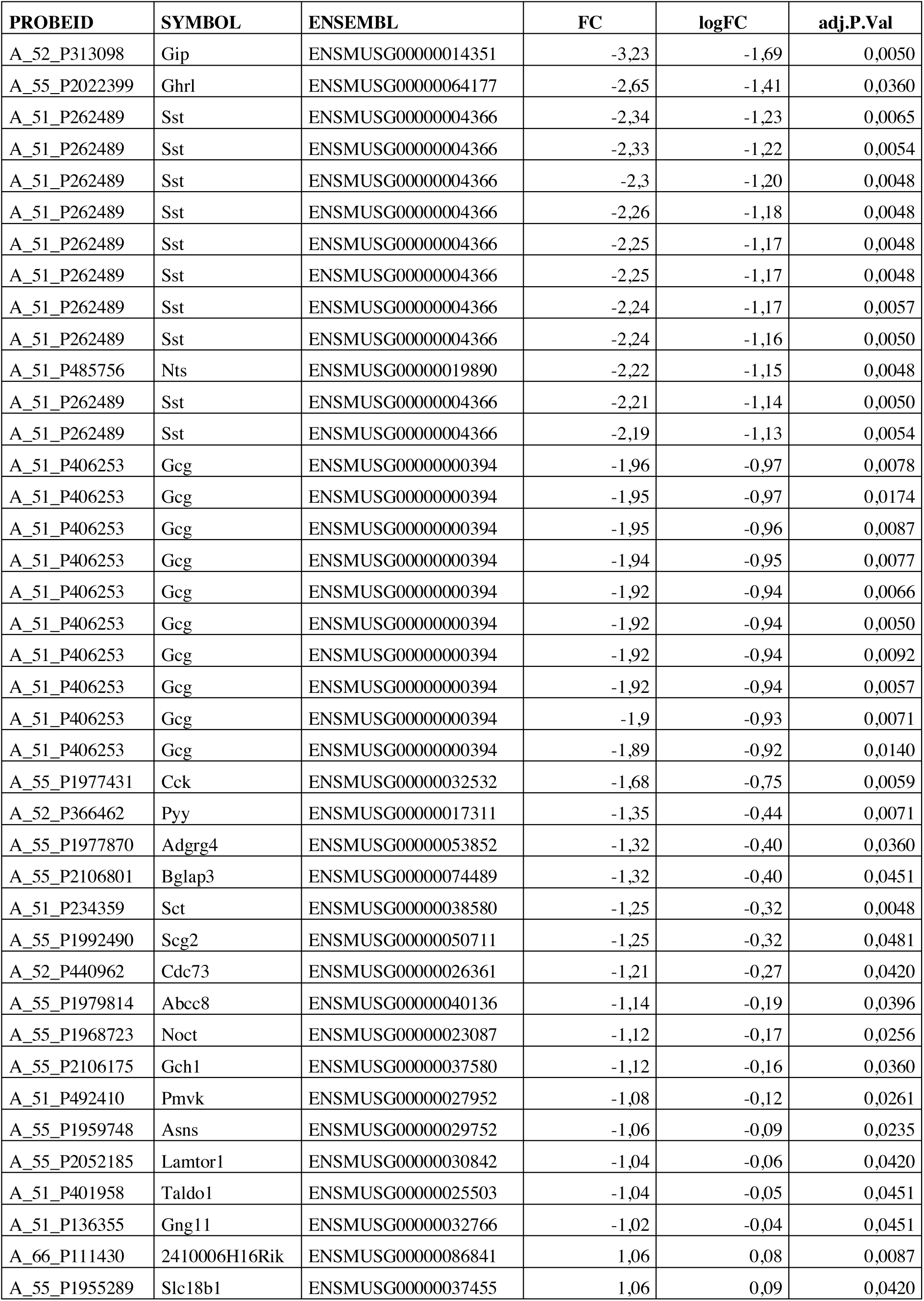
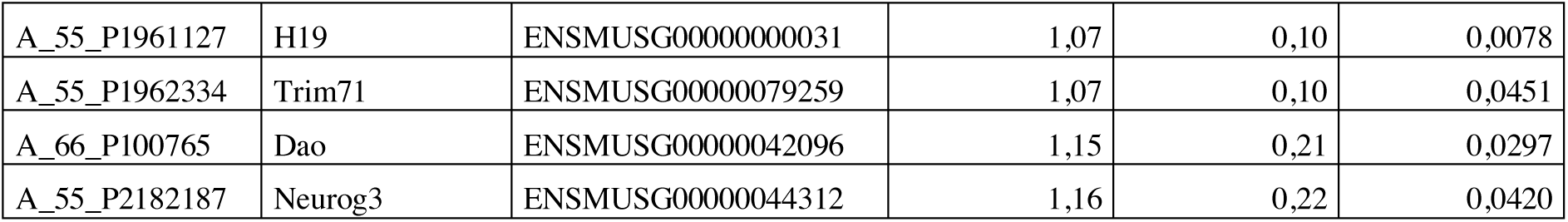
List of genes differentially regulated in Rfx6^ΔEndo^ small intestines (E18.5) compared to controls ordered by Fold Change.

**Table S3 :**
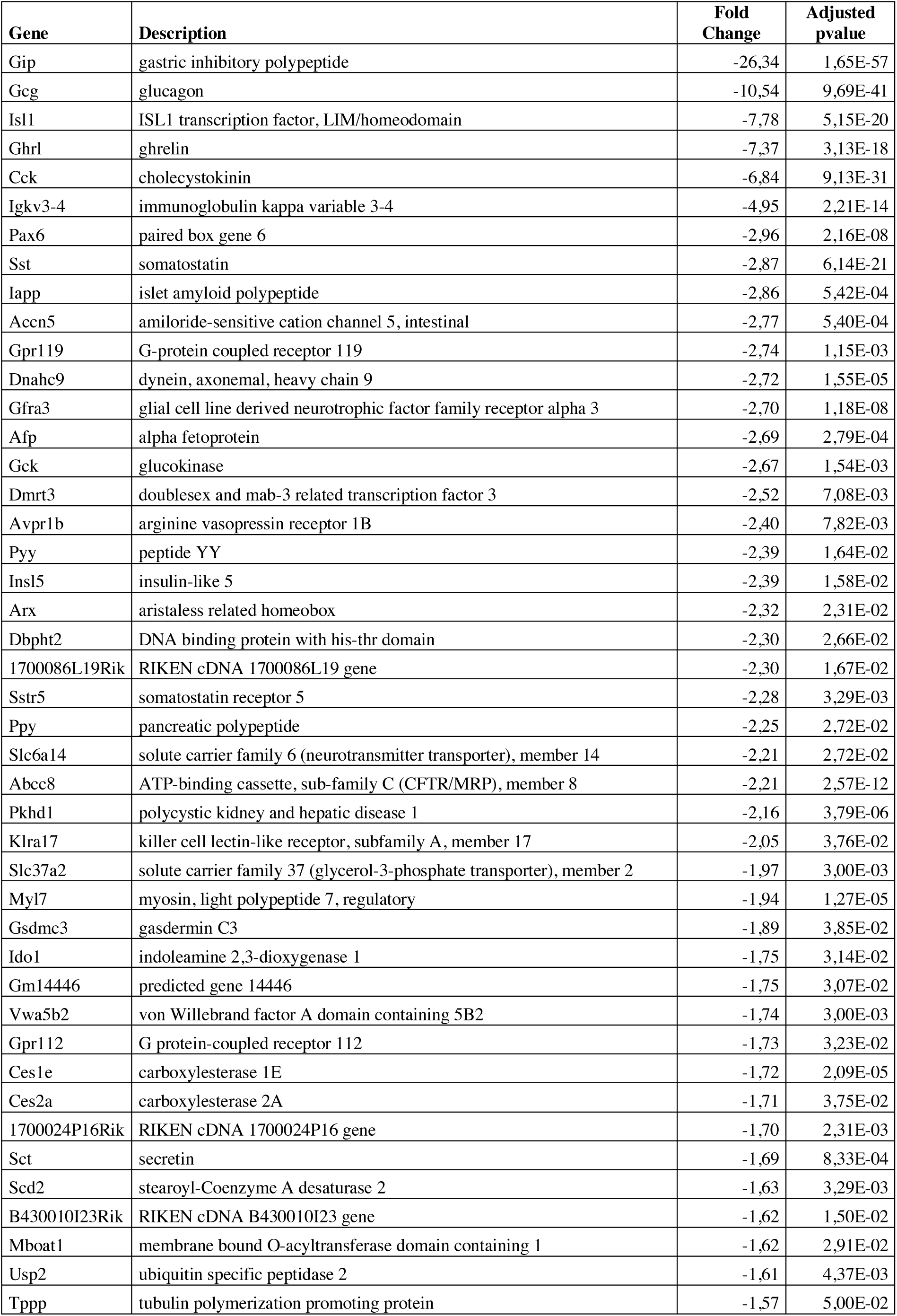
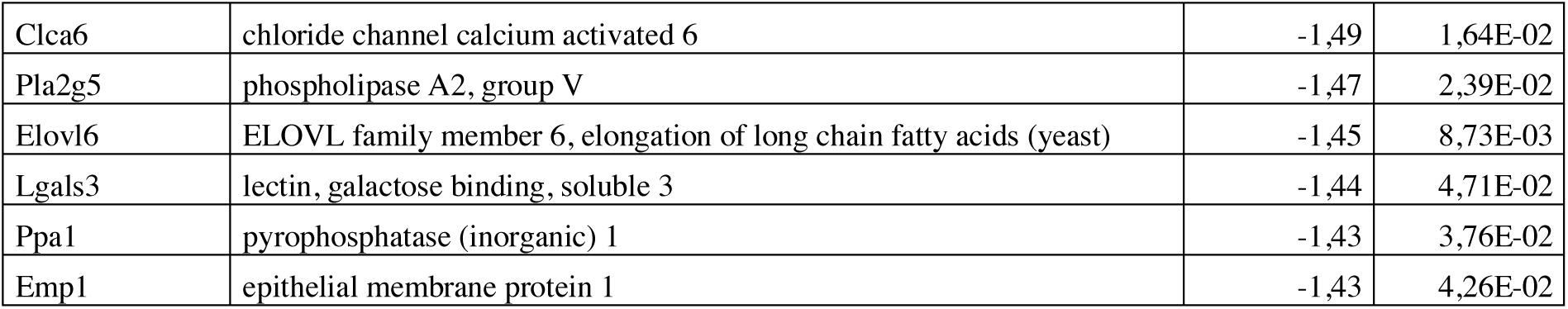
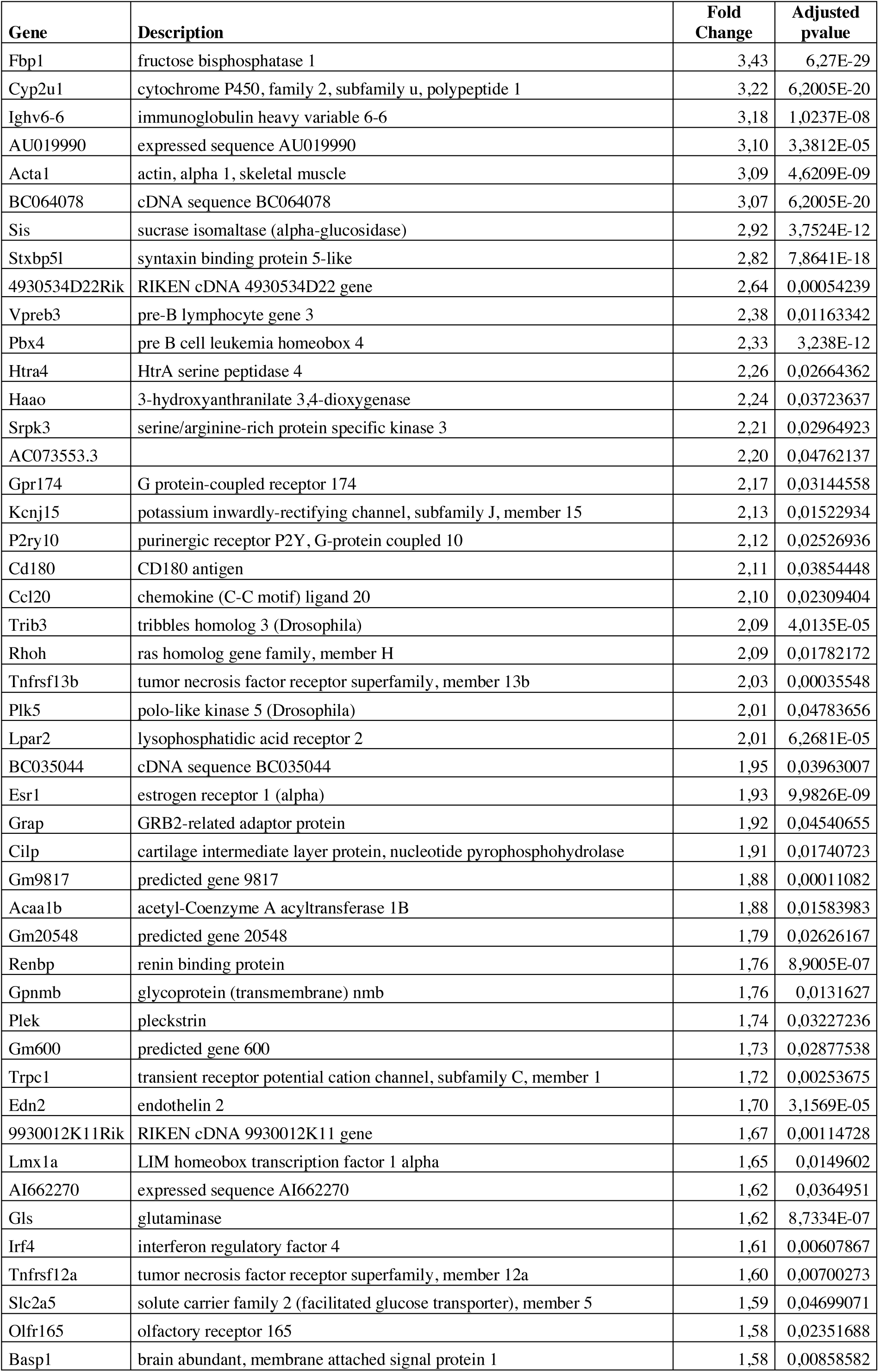
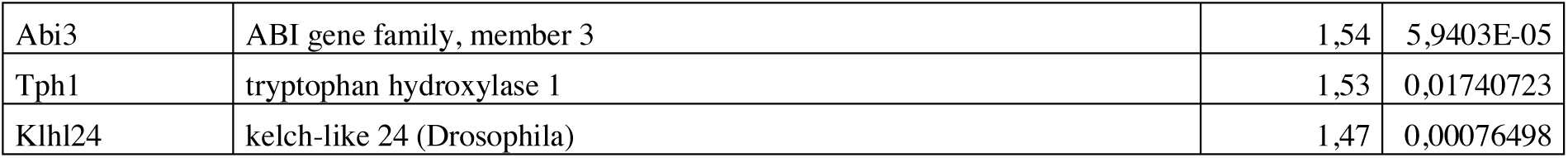
List of 50 genes downregulated or upregulated in the ileum of Rfx6^ΔAdInt^ mice 1week after tamoxifen treatment ordered by fold change.

**Table S4 :**
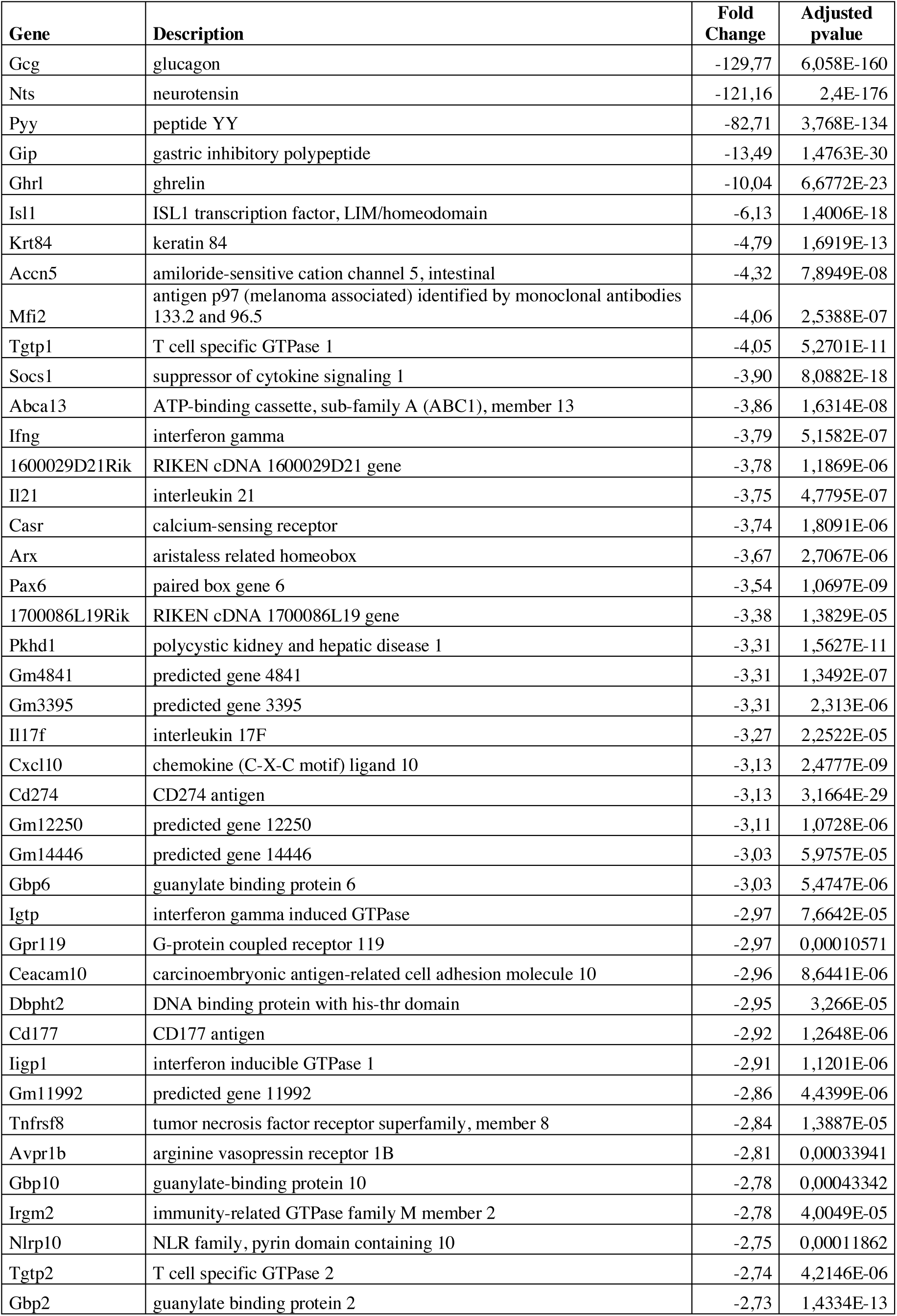
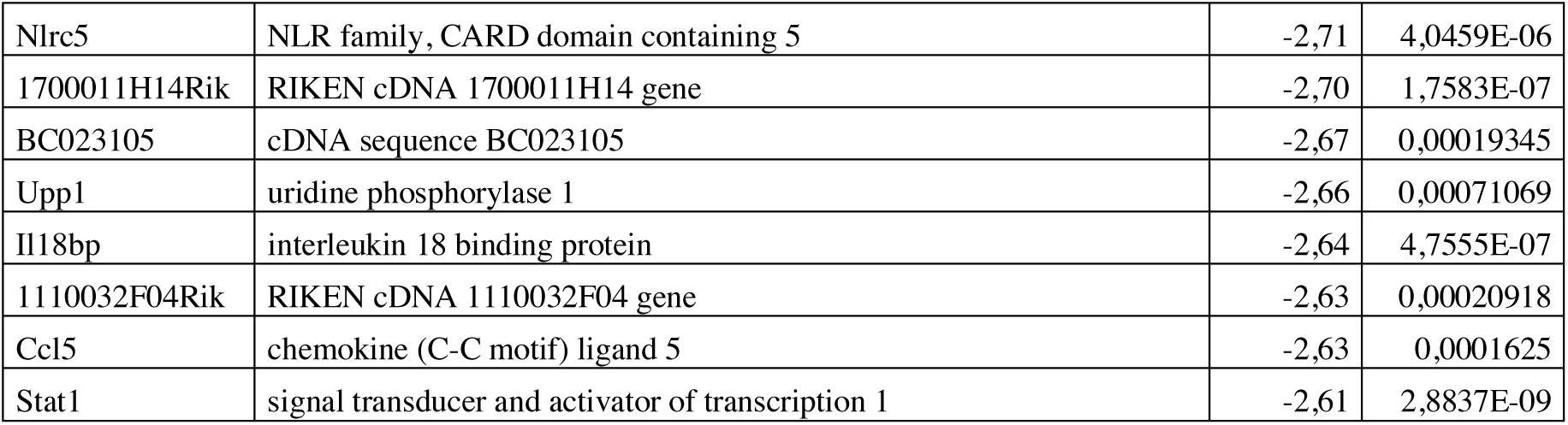
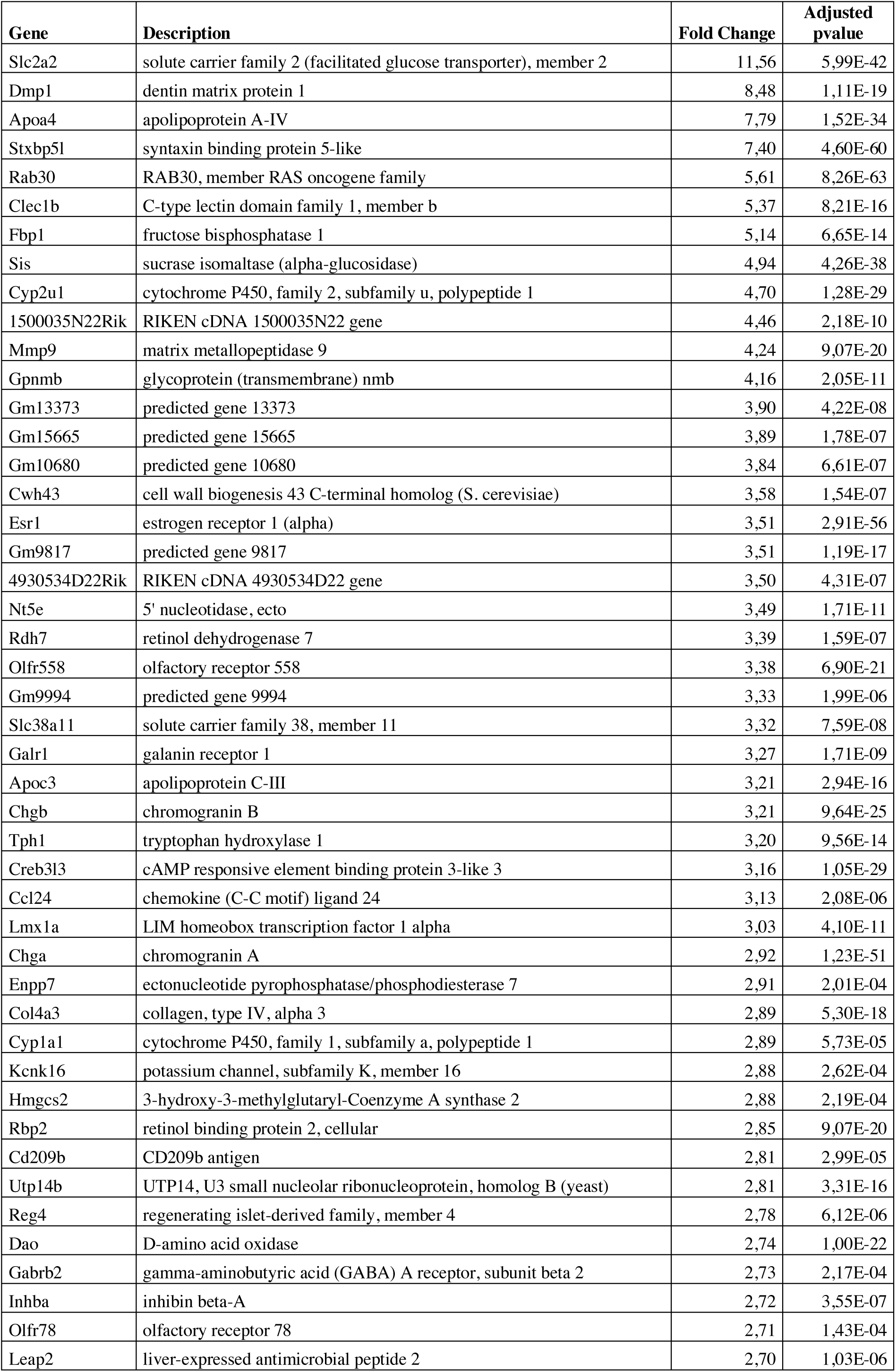
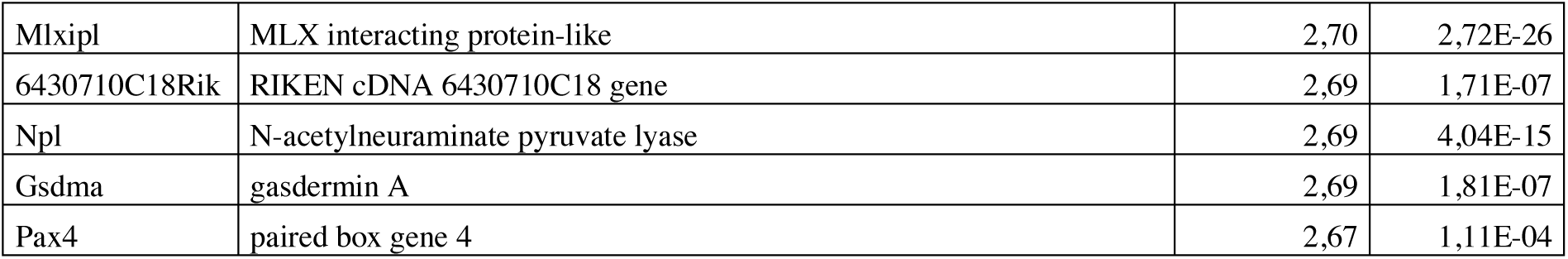
List of 50 genes downregulated or upregulated in the ileum of Rfx6^ΔAdInt^ mice 3 months after tamoxifen treatment ordered by fold change.

## Bibliography

[1] Egerod, K.L., Engelstoft, M.S., Grunddal, K.V., Nøhr, M.K., Secher, A., Sakata, I., et al., 2012. A Major Lineage of Enteroendocrine Cells Coexpress CCK, Secretin, GIP, GLP-1, PYY, and Neurotensin but Not Somatostatin. Endocrinology.

[2] Habib, A.M., Richards, P., Cairns, L.S., Rogers, G.J., Bannon, C.A.M., Parker, H.E., et al., 2012. Overlap of endocrine hormone expression in the mouse intestine revealed by transcriptional profiling and flow cytometry. Endocrinology 153(7):3054–3065.

[3] Grün, D., Lyubimova, A., Kester, L., Wiebrands, K., Basak, O., Sasaki, N., et al., 2015. Single-cell messenger RNA sequencing reveals rare intestinal cell types. Nature 525(7568):251–255.

[4] Haber, A.L., Biton, M., Rogel, N., Herbst, R.H., Shekhar, K., Smillie, C., et al., 2017. A single-cell survey of the small intestinal epithelium. Nature 551(7680):1–28.

[5] Drucker, D.J., 2016. Evolving Concepts and Translational Relevance of Enteroendocrine Cell Biology. Journal of Clinical Endocrinology & Metabolism 101(3):778–786.

[6] Beumer, J., Artegiani, B., Post, Y., Reimann, F., Gribble, F., Nguyen, T.N., et al., 2018. Enteroendocrine cells switch hormone expression along the crypt-to-villus BMP signalling gradient. Nature Cell Biology 20(8):909–916.

[7] Shivdasani, R.A., 2018. Limited gut cell repertoire for multiple hormones. Nature Cell Biology.

[8] Jenny, M., 2002. Neurogenin3 is differentially required for endocrine cell fate specification in the intestinal and gastric epithelium. The EMBO Journal 21(23):6338–6347.

[9] Lee, C.S., Perreault, N., Brestelli, J.E., Kaestner, K.H., 2002. Neurogenin 3 is essential for the proper specification of gastric enteroendocrine cells and the maintenance of gastric epithelial cell identity. Genes & development 16(12):1488–1497.

[10] Schonhoff, S.E., Giel-Moloney, M., Leiter, A.B., 2004. Neurogenin 3-expressing progenitor cells in the gastrointestinal tract differentiate into both endocrine and non-endocrine cell types. 270(2):443–454.

[11] Mellitzer, G., Beucher, A., Lobstein, V., Michel, P., Robine, S., Kedinger, M., et al., 2010. Loss of enteroendocrine cells in mice alters lipid absorption and glucose homeostasis and impairs postnatal survival. The Journal of clinical investigation 120(5):1708–1721.

[12] Shroyer, N.F., Helmrath, M.A., Wang, V.Y.C., Antalffy, B., Henning, S.J., Zoghbi, H.Y., 2007. Intestine-specific ablation of mouse atonal homolog 1 (Math1) reveals a role in cellular homeostasis. YGAST 132(7):2478–2488.

[13] Naya, F.J., Huang, H.P., Qiu, Y., Mutoh, H., DeMayo, F.J., Leiter, A.B., et al., 1997. Diabetes, defective pancreatic morphogenesis, and abnormal enteroendocrine differentiation in BETA2/neuroD-deficient mice. Genes & development 11(18):2323–2334.

[14] Beucher, A., Gjernes, E., Collin, C., Courtney, M., Meunier, A., Collombat, P., et al., 2012. The homeodomain-containing transcription factors Arx and Pax4 control enteroendocrine subtype specification in mice. PLoS One 7(5):e36449.

[15] Du, A., McCracken, K.W., Walp, E.R., Terry, N.A., Klein, T.J., Han, A., et al., 2012. Arx is required for normal enteroendocrine cell development in mice and humans. 365(1):175–188.

[16] Larsson, L.I., St-Onge, L., Hougaard, D.M., Sosa-Pineda, B., Gruss, P., 1998. Pax 4 and 6 regulate gastrointestinal endocrine cell development. Mechanisms of Development 79(1-2):153–159.

[17] Terry, N.A., Walp, E.R., Lee, R.A., Kaestner, K.H., May, C.L., 2014. Impaired enteroendocrine development in intestinal-specific Islet1 mouse mutants causes impaired glucose homeostasis. AJP: Gastrointestinal and Liver Physiology 307(10):G979–G991.

[18] Ye, D.Z., Kaestner, K.H., 2009. Foxa1 and Foxa2 control the differentiation of goblet and enteroendocrine L- and D-cells in mice. Gastroenterology 137(6):2052–2062.

[19] Gierl, M.S., Karoulias, N., Wende, H., Strehle, M., Birchmeier, C., 2006. The zinc-finger factor Insm1 (IA-1) is essential for the development of pancreatic beta cells and intestinal endocrine cells. Genes & development 20(17):2465–2478.

[20] Soyer, J., Flasse, L., Raffelsberger, W., Beucher, A., Orvain, C., Peers, B., et al., 2010. Rfx6 is an Ngn3-dependent winged helix transcription factor required for pancreatic islet cell development. Development (Cambridge, England) 137(2):203–212.

[21] Smith, S.B., Qu, H.-Q., Taleb, N., Kishimoto, N.Y., Scheel, D.W., Lu, Y., et al., 2010. Rfx6 directs islet formation and insulin production in mice and humans. Nature 463(7282):775–780.

[22] Piccand, J., Strasser, P., Hodson, D.J., Meunier, A., Ye, T., Keime, C., et al., 2014. Rfx6 maintains the functional identity of adult pancreatic β cells. Cell reports 9(6):2219–2232.

[23] Suzuki, K., Harada, N., Yamane, S., Nakamura, Y., Sasaki, K., Nasteska, D., et al., 2013. Transcriptional Regulatory Factor X6 (Rfx6) Increases Gastric Inhibitory Polypeptide (GIP) Expression in Enteroendocrine K-cells and Is Involved in GIP Hypersecretion in High Fat Diet-induced Obesity. The Journal of biological chemistry 288(3):1929–1938.

[24] Pearl, E.J., Jarikji, Z., Horb, M.E., 2011. Functional analysis of Rfx6 and mutant variants associated with neonatal diabetes. 351(1):135–145.

[25] Spiegel, R., Dobbie, A., Hartman, C., de Vries, L., Ellard, S., Shalev, S.A., 2011. Clinical characterization of a newly described neonatal diabetes syndrome caused by RFX6 mutations. American journal of medical genetics. Part A 155A(11):2821–2825.

[26] Concepcion, J.P., Reh, C.S., Daniels, M., Liu, X., Paz, V.P., Ye, H., et al., 2014. Neonatal diabetes, gallbladder agenesis, duodenal atresia, and intestinal malrotation caused by a novel homozygous mutation in RFX6. Pediatric diabetes 15(1):67–72.

[27] Artuso, R., Provenzano, A., Mazzinghi, B., Giunti, L., Palazzo, V., Andreucci, E., et al., 2014. Therapeutic implications of novel mutations of the RFX6 gene associated with early-onset diabetes. Pharmacogenomics J.

[28] Zegre Amorim, M., Houghton, J.A.L., Carmo, S., Salva, I., Pita, A., Pereira-da-Silva, L., 2015. Mitchell-Riley Syndrome: A Novel Mutation in RFX6 Gene. Case reports in genetics 2015:937201.

[29] Sansbury, F.H., Kirel, B., Caswell, R., Lango Allen, H., Flanagan, S.E., Hattersley, A.T., et al., 2015. Biallelic RFX6 mutations can cause childhood as well as neonatal onset diabetes mellitus. European journal of human genetics : EJHG 23(12):1744–1748.

[30] Skopkova, M., Ciljakova, M., Havlicekova, Z., Vojtkova, J., Valentinova, L., Danis, D., et al., 2016. Two novel RFX6 variants in siblings with Mitchell-Riley syndrome with later diabetes onset and heterotopic gastric mucosa. European Journal of Medical Genetics 59(9):429–435.

[31] Chandra, V., Albagli-Curiel, O., Hastoy, B., Piccand, J., Randriamampita, C., Vaillant, E., et al., 2014. RFX6 regulates insulin secretion by modulating Ca2+ homeostasis in human β cells. Cell reports 9(6):2206–2218.

[32] Khan, N., Dandan, W., Al Hassani, N., Hadi, S., 2016. A Newly-Discovered Mutation in the RFX6 Gene of the Rare Mitchell-Riley Syndrome. J Clin Res Pediatr Endocrinol 8(2):246–249.

[33] Piccand, J., Strasser, P., Hodson, D.J., Meunier, A., Ye, T., Keime, K., et al., 2014. Rfx6 maintains the functional identity of adult pancreatic β-cells. Manuscript in revision.

[34] Yoshida, S., Takakura, A., Ohbo, K., Abe, K., Wakabayashi, J., Yamamoto, M., et al., 2004. Neurogenin3 delineates the earliest stages of spermatogenesis in the mouse testis., Dev. Biol., pp. 447–458.

[35] El Marjou, F., Janssen, K.-P., Chang, B.H.-J., Li, M., Hindie, V., Chan, L., et al., 2004. Tissue-specific and inducible Cre-mediated recombination in the gut epithelium., Genesis, pp. 186–193.

[36] Mellitzer, G., Martin, M., Sidhoum-Jenny, M., Orvain, C., Barths, J., Seymour, P.A., et al., 2004. Pancreatic islet progenitor cells in neurogenin 3-yellow fluorescent protein knock-add-on mice. Mol.Endocrinol. 18(11):2765–2776.

[37] Bolstad, B.M., Irizarry, R.A., Astrand, M., Speed, T.P., 2003. A comparison of normalization methods for high density oligonucleotide array data based on variance and bias. Bioinformatics 19(2):185–193.

[38] Ritchie, M.E., Phipson, B., Wu, D., Hu, Y., Law, C.W., Shi, W., et al., 2015. limma powers differential expression analyses for RNA-sequencing and microarray studies. Nucleic Acids Res 43(7):e47.

[39] Love, M.I., Huber, W., Anders, S., 2014. Moderated estimation of fold change and dispersion for RNA-seq data with DESeq2. Genome Biol 15(12):550.

[40] Smith, T., Heger, A., Sudbery, I., 2017. UMI-tools: modeling sequencing errors in Unique Molecular Identifiers to improve quantification accuracy. Genome Res 27(3):491–499.

[41] Satija, R., Farrell, J.A., Gennert, D., Schier, A.F., Regev, A., 2015. Spatial reconstruction of single-cell gene expression data. Nat Biotechnol 33(5):495–502.

[42] Robinson, M.D., McCarthy, D.J., Smyth, G.K., 2010. edgeR: a Bioconductor package for differential expression analysis of digital gene expression data. Bioinformatics 26(1):139–140.

[43] Soneson, C., Robinson, M.D., 2018. Bias, robustness and scalability in single-cell differential expression analysis. Nat Methods 15(4):255–261.

[44] Haghverdi, L., Buettner, F., Theis, F.J., 2015. Diffusion maps for high-dimensional single-cell analysis of differentiation data. Bioinformatics 31(18):2989–2998.

[45] Gross, S., Garofalo, D.C., Balderes, D.A., Mastracci, T.L., Dias, J.M., Perlmann, T., et al., 2016. The novel enterochromaffin marker Lmx1a regulates serotonin biosynthesis in enteroendocrine cell lineages downstream of Nkx2.2. Development (Cambridge, England) 143(14):2616–2628.

[46] Croset, M., Rajas, F., Zitoun, C., Hurot, J.M., Montano, S., Mithieux, G., 2001. Rat small intestine is an insulin-sensitive gluconeogenic organ. Diabetes 50(4):740–746.

[47] Kellett, G.L., Brot-Laroche, E., Mace, O.J., Leturque, A., 2008. Sugar absorption in the intestine: the role of GLUT2. Annual review of nutrition 28:35–54.

[48] Lee, A.-H., 2012. The role of CREB-H transcription factor in triglyceride metabolism. Current Opinion in Lipidology 23(2):141–146.

[49] Kim, T.-H., Saadatpour, A., Guo, G., Saxena, M., Cavazza, A., Desai, N., et al., 2016. Single-Cell Transcript Profiles Reveal Multilineage Priming in Early Progenitors Derived from Lgr5+ Intestinal Stem Cells. Cell reports 16(8):2053–2060.

[50] Gehart, H., van Es, J.H., Hamer, K., Beumer, J., Kretzschmar, K., Dekkers, J.F., et al., 2019. Identification of Enteroendocrine Regulators by Real-Time Single-Cell Differentiation Mapping. Cell 176(5):1158–1173 e1116.

[51] Desai, S., Loomis, Z., Pugh-Bernard, A., Schrunk, J., Doyle, M.J., Minic, A., et al., 2008. Nkx2.2 regulates cell fate choice in the enteroendocrine cell lineages of the intestine. 313(1):58–66.

[52] Zeisel, A., Hochgerner, H., Lonnerberg, P., Johnsson, A., Memic, F., van der Zwan, J., et al., 2018. Molecular Architecture of the Mouse Nervous System. Cell 174(4):999–1014 e1022.

[53] Collombat, P., 2003. Opposing actions of Arx and Pax4 in endocrine pancreas development. Genes & development 17(20):2591–2603.

